# Slide-tags: scalable, single-nucleus barcoding for multi-modal spatial genomics

**DOI:** 10.1101/2023.04.01.535228

**Authors:** Andrew J. C. Russell, Jackson A. Weir, Naeem M. Nadaf, Matthew Shabet, Vipin Kumar, Sandeep Kambhampati, Ruth Raichur, Giovanni J. Marrero, Sophia Liu, Karol S. Balderrama, Charles R. Vanderburg, Vignesh Shanmugam, Luyi Tian, Catherine J. Wu, Charles H. Yoon, Evan Z. Macosko, Fei Chen

## Abstract

Recent technological innovations have enabled the high-throughput quantification of gene expression and epigenetic regulation within individual cells, transforming our understanding of how complex tissues are constructed. Missing from these measurements, however, is the ability to routinely and easily spatially localise these profiled cells. We developed a strategy, Slide-tags, in which single nuclei within an intact tissue section are ‘tagged’ with spatial barcode oligonucleotides derived from DNA-barcoded beads with known positions. These tagged nuclei can then be used as input into a wide variety of single-nucleus profiling assays. Application of Slide-tags to the mouse hippocampus positioned nuclei at less than 10 micron spatial resolution, and delivered whole-transcriptome data that was indistinguishable in quality from ordinary snRNA-seq. To demonstrate that Slide-tags can be applied to a wide variety of human tissues, we performed the assay on brain, tonsil, and melanoma. We revealed cell-type-specific spatially varying gene expression across cortical layers and spatially contextualised receptor-ligand interactions driving B-cell maturation in lymphoid tissue. A major benefit of Slide-tags is that it is easily adaptable to virtually any single-cell measurement technology. As proof of principle, we performed multiomic measurements of open chromatin, RNA, and T-cell receptor sequences in the same cells from metastatic melanoma. We identified spatially distinct tumour subpopulations to be differentially infiltrated by an expanded T-cell clone and undergoing cell state transition driven by spatially clustered accessible transcription factor motifs. Slide-tags offers a universal platform for importing the compendium of established single-cell measurements into the spatial genomics repertoire.

## Introduction

Technology development efforts in genomics during the last decade have produced an extensive toolkit of single-cell and single-nucleus sequencing methods, enabling high-throughput molecular characterization of many macromolecules^1–7^. Missing from these measurements, however, is the cytoarchitectural organisation of the cells being profiled. Spatially-resolved sequencing technologies aim to address this drawback by barcoding macromolecules with oligonucleotides whose spatial positions are known^8–12^. However, direct transfer of design principles from single-cell sequencing methods to spatially-resolved profiling is often impossible, necessitating the re-invention of each molecular assay (e.g. transcriptomics^9, 10^, mutations^8^, or ATAC-seq^13–15^) in a spatial context. Furthermore, while single-cell computational tools are extremely mature^16^, additional sources of noise in spatial genomics techniques require their re-design as well, for example to address problems with cellular mixing^17–19^. An alternative to capture-based strategies is to isolate single cells while retaining spatial barcoding information; so far, this has been demonstrated only at limited spatial resolution, and with sparse sampling of tissues^20, 21^. An ideal spatial genomics technology would: 1) efficiently capture cell profiles from tissue sections; 2) resolve cellular positions at low-micrometre resolution; and 3) be generally applicable to any single-cell methodology.

Here we introduce Slide-tags, a method in which cellular nuclei from an intact fresh frozen tissue section are ‘tagged’ with spatial barcode oligonucleotides derived from DNA-barcoded beads with known positions. Isolated nuclei are then profiled with existing single-cell methods with the addition of spatial positions. We demonstrate the tissue versatility of Slide-tags by assaying adult and developing mouse brain, human cerebral cortex, human tonsil, and human melanoma. Across tissues and species, we import spatially “tagged” nuclei into standard workflows for single-nucleus RNA-seq (snRNA-seq), single-nucleus ATAC-seq (snATAC-seq), and T-cell receptor (TCR) sequencing. Slide-tags is also readily integrated into established single-cell computational workflows, such as copy number variation (CNV) inference. In doing so, we leverage the truly single-cell, spatially resolved, multimodal capacity of Slide-tags to reveal cell-type specific spatially varying gene expression, spatially contextualise receptor-ligand interactions, and dissect genetic and epigenetic factors participating in tumour microenvironments.

## Results

### Labelling of nuclei with spatial oligonucleotide barcodes

We previously developed densely packed spatially indexed arrays of DNA-barcoded 10 μm beads, generated using split-pool phosphoramidite synthesis and indexed by sequencing-by-ligation^8, 10, 22^. In our original Slide-seq methodology, DNA or RNA from tissues is captured and spatially barcoded using these arrays. In our new Slide-tags technology, we photocleave and diffuse these bead-derived spatial barcodes into 20 μm fresh frozen tissue sections to associate them with nuclei (Fig. 1a). We postulated that once these barcodes are associated with nuclei, they could be used as input to established single nucleus sequencing approaches (Methods) with only minor protocol modifications.

**Fig. 1:**
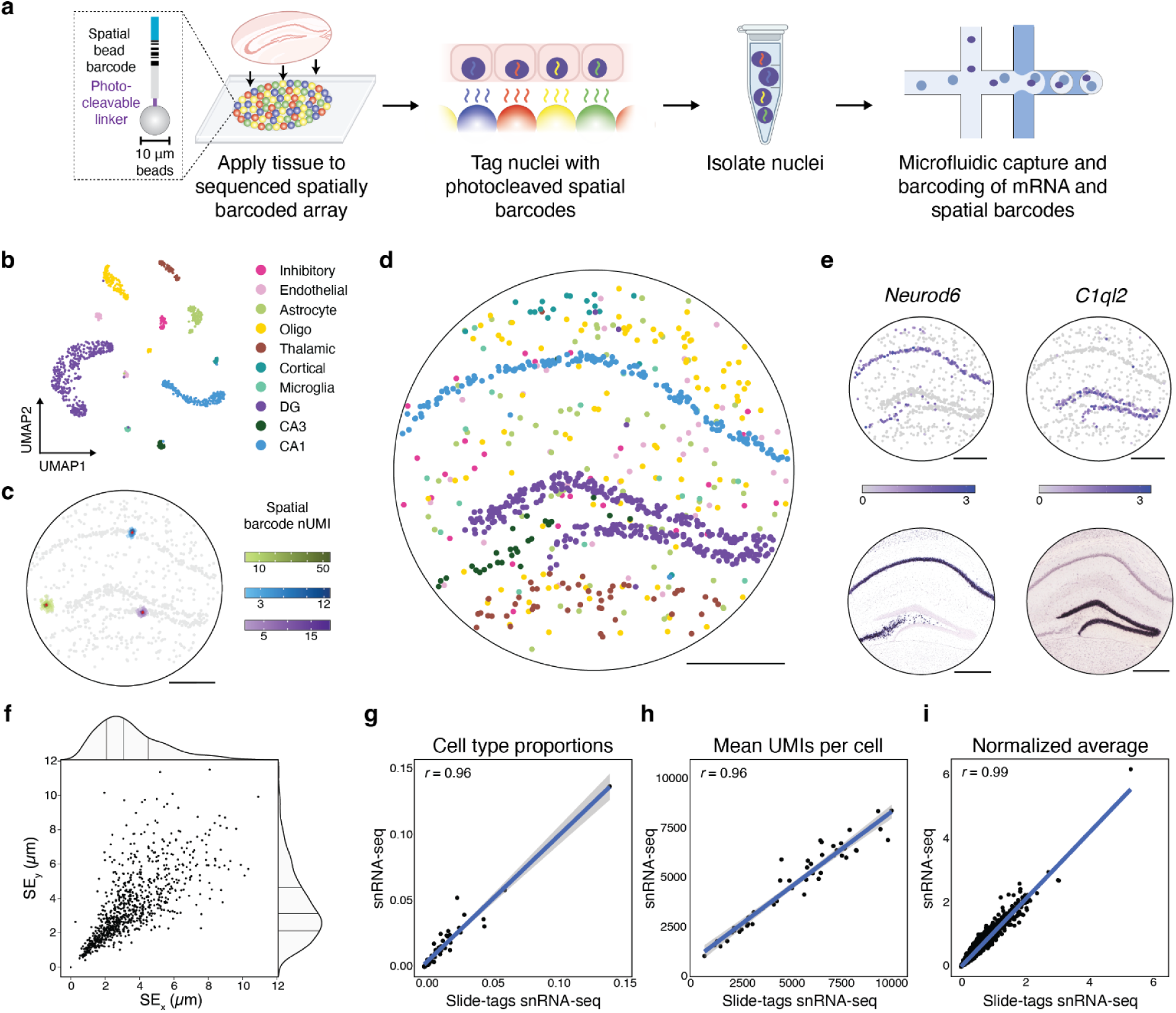
Slide-tags enables single-nucleus spatial transcriptomics in mouse hippocampus. **a,** Schematic of Slide-tags. A 20 μm fresh-frozen tissue section is applied to a monolayer of randomly deposited, DNA barcoded beads that have been spatially indexed. These DNA spatial barcodes are photocleaved, and associate with nuclei. Spatially barcoded nuclei are then profiled using established droplet-based single-nucleus sequencing technologies. **b,** UMAP embedding of snRNA-seq profiles coloured by cell-type annotations. **c,** Plots showing the signal spatial barcode clusters for select cells, coloured by cell-type annotations and number of spatial barcode UMIs **d,** Slide-tags applied to mouse hippocampus enables localization of single nuclei to spatial coordinates, cells are coloured by cell-type annotation as in b. **e,** Spatial expression of known marker genes compared to in situ hybridization data from Allen Mouse Brain Atlas^25^. **f,** Plot showing the standard error (SE) for each singlet true spatial barcode cluster in x and y. Density shows: centre line, median; adjacent lines, upper and lower quartiles; **g-i,** Comparison metrics plotted for snRNA-seq compared to Slide-tags snRNA-seq, performed on consecutive sections. 95% Confidence intervals are plotted. **g,** Cell type proportions and mean UMIs per cell is plotted by cell type. **h,** Normalised average count is per gene across all cells. **i,** Expression counts for each cell were divided by the total counts for that cell and multiplied by 10,000, this value + 1 is then natural-log transformed. *r* is the Pearson correlation coefficient. All scale bars denote 500 μm. DG = dentate gyrus, Oligo = oligodendrocyte, CA1 = Cornu Ammonis area 1, CA3 = Cornu Ammonis area 3.

### Spatially-resolved single-nucleus RNA sequencing of the mouse brain

To benchmark our approach, we performed Slide-tags followed by droplet-based snRNA-seq on a 20 μm coronal section of the adult mouse hippocampus, which has a highly stereotyped architecture that is useful for validating spatial techniques^10^. We dissociated and sequenced 1,661 nuclei from a 3 mm^2^ area coronal tissue section, clustering the data using a standard single cell pipeline^23^ (Fig. 1b), and annotating clusters using well-established cell class markers (Fig. S1). Multiple spatial barcodes were detected per nucleus, allowing higher assignment confidence than protocols where only one spatial barcode is associated with a cell (Fig. 1c). To spatially position our single nucleus transcriptomes, we utilised density-based spatial clustering of applications with noise (DBSCAN)^24^ to separate background spatial barcodes from true signal (Fig. S2, Methods). Nuclei are then assigned a spatial coordinate using the UMI-weighted centroid of the DBSCAN-clustered spatial barcodes denoted true signal (Methods). Using this procedure, we assigned spatial locations to 839 high-quality nuclei profiles (50.5% of profiled nuclei, 11250 median UMIs per nucleus). Examination of the spatial positions of individual clusters recapitulated the expected cytoarchitectural arrangement of the hippocampus (Fig. 1d). Furthermore, spatial expression profiles of individual genes matched existing in situ hybridization data^25^ (Fig. 1e). To quantify spatial positioning accuracy, we first compared the width of the hippocampal subfield CA1 in Slide-tags with a Nissl-stained serial section and found the width of the Slide-tags feature was congruent with the Nissl image (Fig. S3). Second, we calculated the standard error for each centroid in x and y, and estimated accuracy to be 3.5 ± 1.9 μm in x and 3.5 ± 2 μm in y (mean ± s.d., Fig. 1f). Third, we quantified the nuclei misassignment rate by leveraging the stereotyped structure of the CA1 and dentate gyrus (DG). We found that 98.7% of CA1 (155/157) and dentate granule (312/316) neurons were localised in the CA1 pyramidal layer, and the DG, respectively (Fig. S1b). We investigated whether the tagging procedure affected resultant snRNA-seq data quality by comparing standard snRNA-seq with Slide-tags followed by snRNA-seq on adjacent sections of mouse hippocampus. We found that recovered cell type proportions (Pearson’s *r* = 0.96, *p* value < 2.2×10^-16^), UMIs recovered per cell (Pearson’s *r* = 0.96, *p* value < 2.2×10^-16^), and gene expression (Pearson’s *r* = 0.99, *p* value < 2.2×10^-16^) were all unaffected by the tagging procedure (Fig. 1g-i). Thus, Slide-tags generated data virtually indistinguishable from snRNA-seq with a theoretical ∼3 micrometre spatial localization accuracy.

We next performed Slide-tags snRNA-seq on a 7 mm^2^ area sagittal section of the embryonic mouse brain at E14 (Fig. S4 a,b), which has been frequently used for benchmarking of new spatial transcriptomics technologies^11, 22, 26^. We sequenced and spatially positioned 4,584 nuclei (4594 median UMIs per nucleus), which we clustered and annotated by cell type (Fig S4 c-e). Compared with existing approaches for single-cell spatial placement, Slide-tags achieved 20-50-fold higher spatial resolution and recovered 4.5-fold more nuclei per unit area. We also recovered 1.8-fold more UMIs and 1.7-fold more genes per nucleus than adjacent technologies at a sequencing saturation of 48% (Fig. S4f).

### Slide-tags enables the identification of layer-specific gene expression across cell types in the human prefrontal cortex

The human cerebral cortex has a well-characterised cytoarchitecture in which specific subpopulations of neurons are arranged in discrete layers. Existing spatial sequencing approaches can resolve broad patterns of spatially varying gene expression in human cortex^27^, but assignment of spatially variable genes to specific cell types is challenging with these methods. We reasoned that Slide-tags could be used for facile profiling of human brain tissue, most especially to discover cell-type-specific spatial gene expression patterns. We profiled a 28.3 mm^2^ region of the human prefrontal cortex from a 78-year-old neurotypical donor (Methods), recovering 4,067 high-quality spatially mapped nuclei with a median of 3,024 UMIs per nucleus (Fig. 2a). Clustering analysis revealed the expected neuronal and glial cell types, recapitulating known layer distributions and spatial structures (Fig. 2b-d, S5a-b). We computationally integrated (Methods) an existing snRNA-seq dataset^28^ that includes layer annotations for 91 neuron subtypes, recovering the expected spatial distributions across subtypes (Fig. 2e-f, S6). Similarly, astrocytes could be clustered into two distinct populations that spatially segregated between white and grey matter regions (Fig. 2g). Quantification of the laminar position of each of these excitatory, inhibitory, and astrocytic populations showed them to be accurately positioned within the white matter and cortical layers (Fig. 2h).

**Fig. 2:**
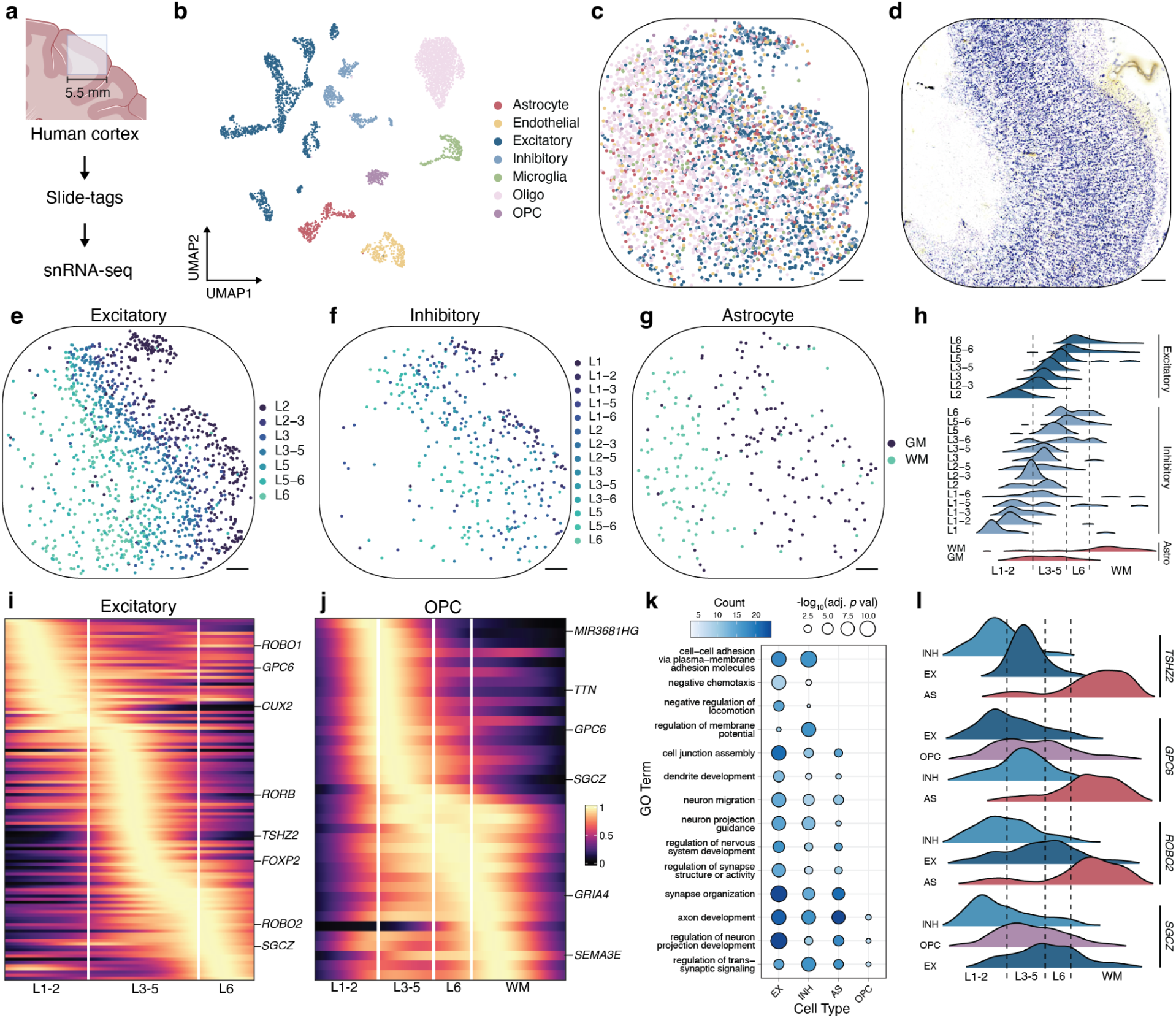
Spatially-resolved cell type-specific expression in the human brain using Slide-tags snRNA-seq. **a,** Schematic of Slide-tags snRNA-seq on a 5.5 mm square region of human prefrontal cortex. **b**, UMAP embedding of snRNA-seq profiles, coloured by cell type assignment. **c**, Spatial mapping of snRNA-seq profiles, coloured by cell type as in b. **d**, A Nissl-stained tissue section adjacent to the profiled section. **e**, Spatial mapping of grouped excitatory neuron subtypes. **f**, Spatial mapping of grouped inhibitory neuron subtypes. **g**, Spatial mapping of white and grey matter astrocytes. **h**, Ridge plot showing the layer specificity of grouped excitatory neuron, grouped inhibitory neuron, and astrocyte subtypes. **i**,**j** Heatmaps of 1D gene expression for excitatory neurons (**i**), and OPCs (**j**). **k**, Gene ontology analysis of the highest spatially variable genes in each cell type. **l**, Ridge plot showing the spatial expression of genes with contrasting gradients across cell types. All scale bars denote 500 μm. Oligo = Oligodendrocyte, OPC = Oligodendrocyte precursor cell, Astro = Astrocyte, INH = Inhibitory, EX = Excitatory, GM = Grey matter, WM = White matter.

We next used our whole-transcriptome, spatially resolved snRNA-seq profiles to systematically identify spatially varying genes in each cell type. We plotted the layer distributions of the highest spatially varying genes (Methods, Table S1) for excitatory neurons (Fig. 2i, S7a), recovering many well-known laminar markers such as *CUX2*, *RORB*, and *FOXP2* (Fig. S5c), as well as for inhibitory neurons (Fig. S5d, S7b) and astrocytes (Fig. S5e, S8a). Interestingly, we also identified spatially varying genes within oligodendrocyte precursor cells (OPCs) which had not been previously known to have areal specialisations (Fig. 2j, S8b). Gene ontology analysis on these spatially varying genes revealed a relationship to biological processes that included cell-cell adhesion, cell junction assembly, and axon development (Fig. 2k, S9, Table S2).

Genes can show spatially variable expression which may derive from several cell types, but assigning such expression variability to individual cell types can be very challenging with traditional spatial transcriptomics approaches because of the mixing of individual pixels. Amongst our spatially varying genes, we identified several that were variable across multiple cell types, such as *SGCZ*, whose spatial expression variation in excitatory and inhibitory neurons was anticorrelated, and showed an orthogonal spatial distribution in OPCs (Fig. 2l). Together, these results demonstrate the ability of Slide-tags to systematically uncover transcriptional variation within the cytoarchitecturally complex tissues of the human brain.

### Spatially-informed receptor-ligand prediction in immune cell-dense human tonsil

A key challenge for spatial genomics technologies is the proper segmentation of densely packed tissues, such as those of immune origin. We reasoned that Slide-tags would be ideal in this setting, given that segmentation is accomplished automatically by dissociating the tissue into individual nuclei. We therefore performed Slide-tags snRNA-seq on human tonsil (Fig. 3a-d), recovering 81,000 nuclei after dissociation from 7 mm^2^ of tissue. We sequenced 8,747 of these nuclei, spatially mapping 5,778 high quality snRNA-seq profiles (2,377 median UMIs per nucleus and 1,557 median genes per nucleus). Clustering of the data identified subpopulations of B and T cells, some of which are known to be spatially segregated (Fig. S10a-b). Indeed, examination of the spatial positions of these clusters revealed the expected spatial architecture of the tissue, with B and T cell zones, as well as germinal centres composed of germinal centre B cells (GCBs), T follicular helper cells, and follicular dendritic cells (Fig. 3c-d, S10b). Sub-classification of GCBs into light zone and dark zone GCBs is challenging using snRNA-seq data alone, as variation in gene expression space is low, requiring many cells to be sampled to uncover the distinction^29^. However, since reactive germinal centres are spatially polarised into light zones and dark zones, we reasoned that we could classify GCBs by harnessing the combined spatial and single-cell data. To do so, we computed spatially varying genes within GCBs via spatial permutation testing^22^, identifying key markers of light zone and dark zone GCBs (Fig. 3e-f, Table S3). Dark zone marker genes included *CXCR4* (Double sided permutation test, Z-score = 7.6, *p* value < 0.001) and *AICDA* (Z-score = 6.9, *p* value < 0.001), genes for dark zone organisation and somatic hypermutation, respectively^30–32^. Light zone-enriched genes included *BCL2A1* (Z-score = 9.1, *p* value < 0.001), an apoptosis regulator gene^33^, and *LMO2* (Z-score = 21.3, *p* value < 0.001), a transcription factor^34, 35^. A subset of expected light zone and dark zone markers had relatively low variance in gene expression, but high spatial permutation effect sizes, demonstrating that spatial positions enhance interpretation of transcriptomic profiles (Fig. S10c, Table S4). Re-clustering GCBs on spatially varying genes enabled classification into dark zone, light zone, and transitional cell states (Fig. 3g, Methods). We then segmented the two largest profiled germinal centres into light zones and dark zones via spatial clustering of dark zone GCBs, the most abundant GCB cell state (Fig. S10d). In corroboration of our zone segmentation, we found T follicular helper cells were enriched in light zones while follicular dendritic cells were dispersed between the light zone and the dark zone (Fig. 3h, Chi-squared_Tfh_ = 43.7, *p* value_Tfh_ = 3.7×10^-11^, Chi-squared_FDC_ = 0.58, *p* value_FDC_ = 0.45).

**Fig. 3:**
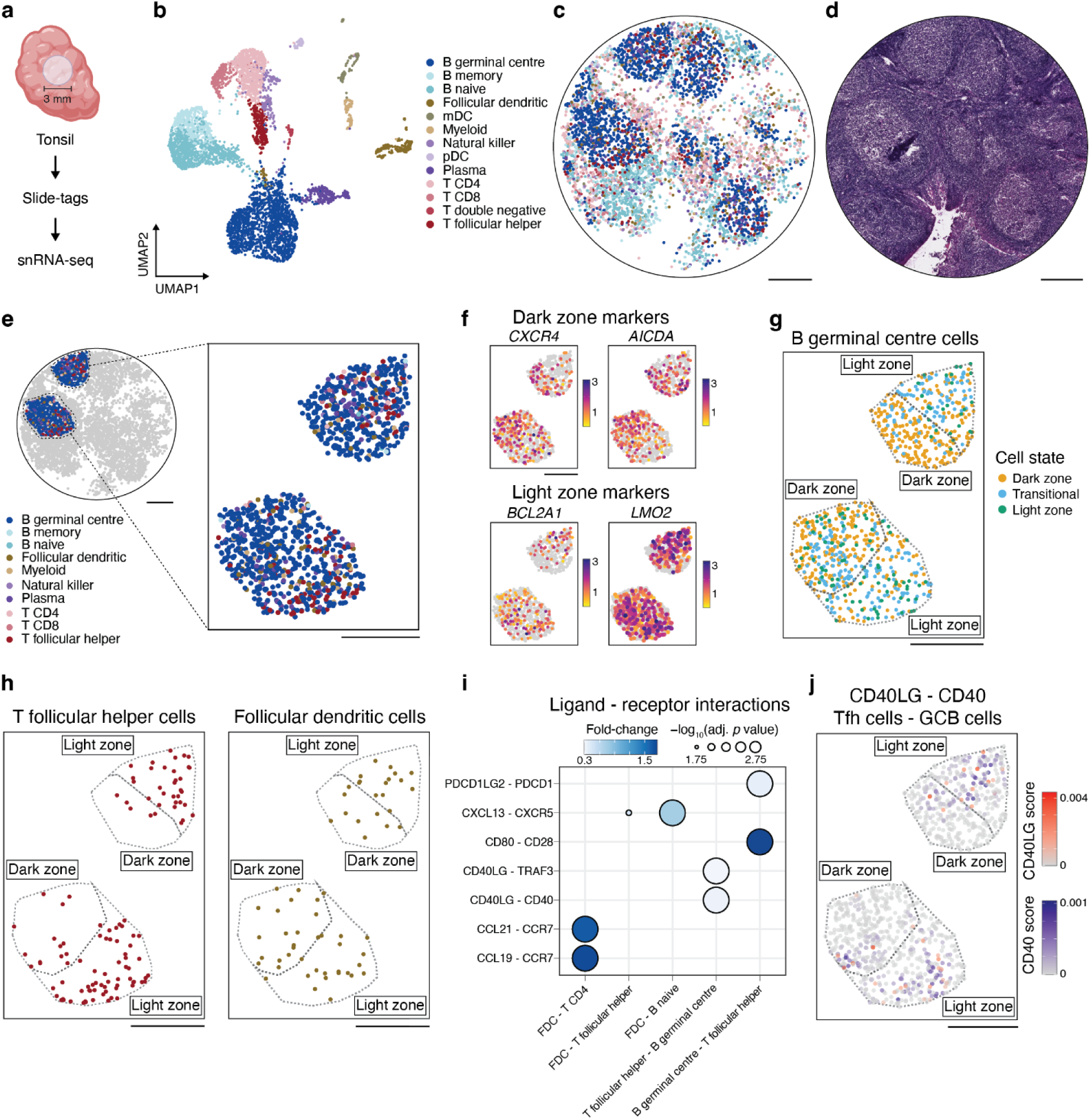
Slide-tags enables cell-type specific spatially varying gene expression analysis and spatial receptor-ligand interaction prediction within human tonsil. **a,** Schematic of Slide-tags snRNA-seq on a 3 mm diameter region of human tonsil tissue. **b,** UMAP embedding of snRNA-seq profiles coloured by cell-type annotations. **c,** Spatial mapping of snRNA-seq profiles, coloured by cell type as in b. **d,** Adjacent H&E-stained section of the profiled region. **e,** Inset of two germinal centres coloured by cell type f, Expression of dark zone and light zone marker genes identified as spatially varying within germinal centres. g, GCB cell state classification and zone segmentation by cluster density of dark zone GCBs. h, Spatial mapping of T follicular helper cells and follicular dendritic cells on zoned germinal centres. i, Dot plot showing select spatially co-occurring receptor-ligand interactions, within certain sender-receiver cell type pairs. j, Spatial mapping of interaction intensity scores for CD40 in GCBs and CD40LG in T follicular helper cells. All scale bars denote 500 μm. mDC = myeloid dendritic cells, pDC = plasmacytoid dendritic cells, GCB = germinal centre B cells, Tfh = T follicular helper cells.

Immune cells engage in extensive cross talk within and around germinal centres^36^. We wondered whether Slide-tags could uncover receptor-ligand interactions that drive such intercellular communication. We first nominated putative receptor-ligand interactions in a spatially agnostic manner using LIANA^37^. We then incorporated spatial information by performing a spatial permutation test to identify interactions that significantly co-occur spatially (Methods). Using this approach, we predicted 645 receptor-ligand interactions, many of which are well-characterised axes of communication during B cell maturation (Fig. 3i, Table S5). For example, we predicted interaction between CD40 and CD40LG within GCBs and T follicular helper cells, a fundamental driver of the germinal centre reaction^38^. We also identified downstream targets of canonical receptor-ligand interactions, such as TRAF3, important in regulating the intracellular effects of CD40-CD40LG binding^39^.

Finally, we reasoned we could spatially contextualise receptor-ligand interactions within native tissue niches. Our predicted interactions can be decomposed into interaction intensity scores for individual cells based on expression and spatial co-occurrence of the receptor and ligand. For the 99 nominated receptor-ligand pairs between GCBs, FDCs, and T follicular helper cells, we used our germinal centre zone segmentations to assess light zone and dark zone enrichment in predicted interaction intensity. We revealed light zone enrichment of 11 interactions and dark zone enrichment of 9 interactions (Fig. S10e, Table S6). GCB CD40 receptor in interaction with T follicular helper cell CD40LG was highly enriched in light zones (Fig. 3j, Wilcoxon rank-sum test, log_2_FC = 1.6, adjusted *p* value = 1.6×10^-9^), while CD40 receptor expression alone was modestly dark zone-biassed (Wilcoxon rank-sum test, log_2_FC = -0.04, *p* value = 0.047). We also revealed zone-biassed interactions with lesser-known importance in the germinal centre reaction, such as the light zone-enriched interaction between T follicular helper cell CD40LG and GCB CD53 (Fig S10e, Wilcoxon rank-sum test, log_2_FC = 1.6, *p* value = 2.3×10^-23^). Altogether, Slide-tags enabled spatial contextualization of cell-type specific receptor-ligand interactions not obvious by analysis of expression alone.

### Slide-tags enables multimodal spatial investigation of metastatic melanoma clones

Epigenetic dysregulation in cancer facilitates drug resistance and pro-metastatic cell state transitions^40–43^. Numerous studies of tumour heterogeneity have revealed clone-specific niches and immune compartments^8, 44, 45^, but the role of epigenetic regulation in establishing and maintaining these spatial niches remains difficult to study. Concurrent spatial mapping of the genome, transcriptome, and epigenomic landscape of the tumour microenvironment could unravel new insights into the complex mechanisms of tumour evolution. Therefore, we developed Slide-tags multiome, enabling simultaneous single-cell spatial profiling of mRNA and chromatin accessibility, along with copy number variation inference.

We first performed Slide-tags snRNA-seq on a metastatic melanoma sample (Fig. S11a-f). We recovered 10,960 nuclei after dissociation from 7 mm^2^ of tissue, sequencing 6,464 of these nuclei and spatially mapping 4,804 high-quality snRNA-seq profiles (2,110 median UMIs per nucleus and 1,317 median genes per nucleus). In an adjacent section, we applied Slide-tags multiome, profiling the tagged nuclei with droplet-based combinatorial snATAC and snRNA-seq (Fig. 4a-b). We spatially mapped 2,529 nuclei from a 38.3 mm^2^ section and both modalities displayed high quality on basic technical performance metrics (Fig. 4b-c, Fig. S12a-e, median UMIs/nucleus = 5,228, median genes/nucleus = 2,429, TSS enrichment score = 11.5, median fragments/nucleus = 1,159, median fraction of unique fragments in peaks = 36.7%).

**Fig. 4:**
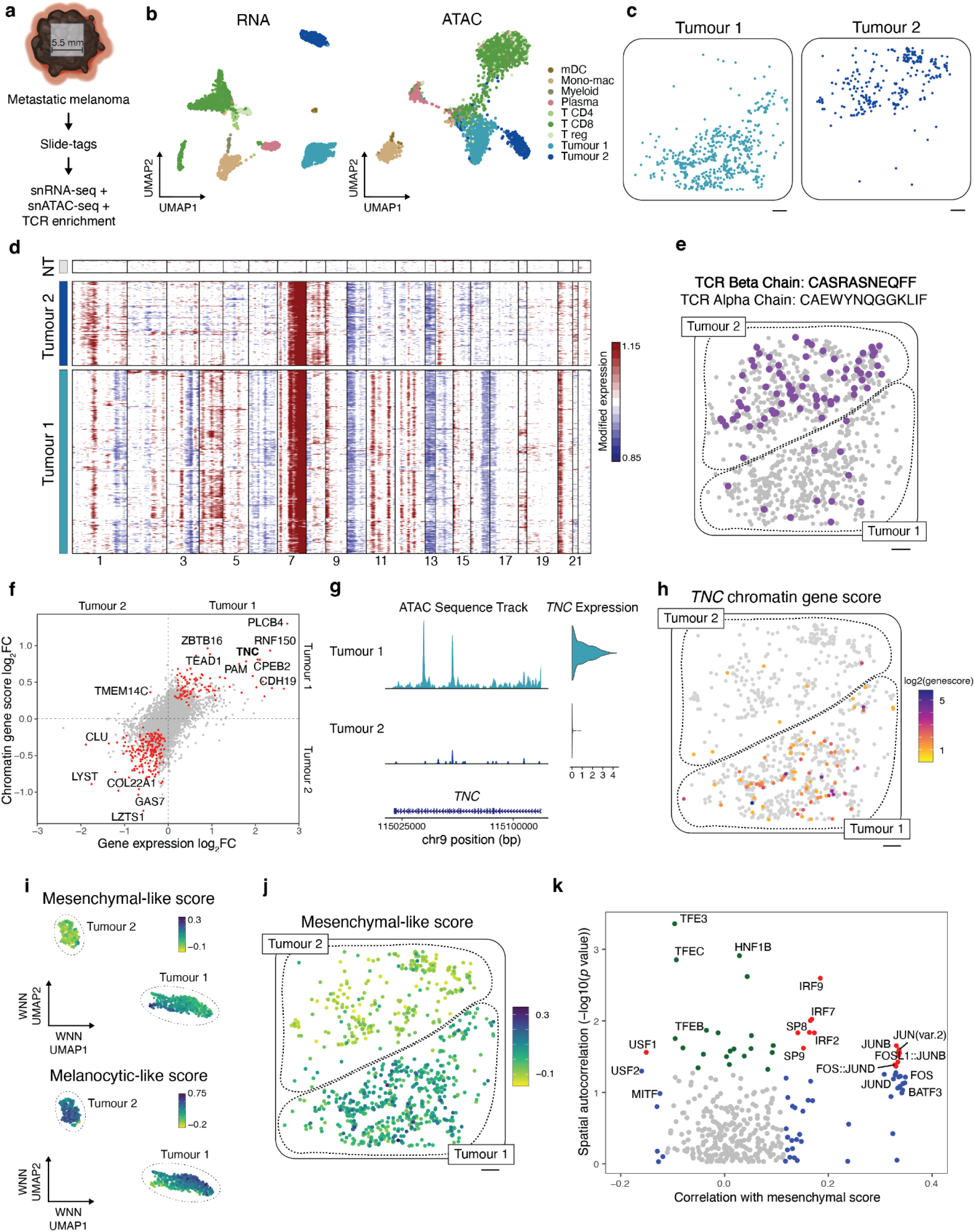
Multiomic Slide-tags captures spatially-resolved clonal relationships between single nuclei in human melanoma. **a,** Schematic representation of joint snATAC-seq and snRNA-seqon a 5.5 mm square region of a human melanoma lymph node metastasis. **b,** UMAP embeddings of snRNA-seq and snATAC-seq profiles coloured by cell type. **c,** Spatial mapping of tumour cluster 1 and tumour cluster 2. **d,** Inferred copy number alterations from transcriptomic data. NT indicates a representative subset of non-tumour cells. **e,** Spatial mapping of a TCR beta chain clonotype expanded in the tumour cluster 2 compartment, with matched alpha chain indicated above. Grey cells depict positions of all CD8 T cells. **f,** Differential gene expression and differential chromatin gene scores between tumour cluster 1 and tumour cluster 2. Red points have adjusted *p* value < 0.05 for both tests. **g,** Genome coverage track and gene expression violin plot of *TNC* between tumour clusters. Range of normalised chromatin accessibility signal is 0-50. **h,** Spatial distribution of *TNC* chromatin accessibility gene scores. Gene scores are log_2_-transformed. **i,** Weighted-nearest neighbours UMAP embedding of tumour cells, with cells coloured by Mesenchymal-like and melanocytic-like cell state scores. **j,** Spatial mapping of mesenchymal-like cell state scores in tumour cells. **k,** Spatial autocorrelation of accessibility in ChromVAR transcription factor motifs correlated with mesenchymal-like cell state scores. Red indicates spatial autocorrelation Moran’s I raw *p* value < 0.05 and significant correlation with mesenchymal-like score (adjusted *p* value < 0.05). Green indicates only spatial autocorrelation raw *p* value < 0.05. Blue indicates only significant correlation with mesenchymal-like score (adj. *p* value < 0.05). Only ChromVAR transcription factor motifs with positive Moran’s I are shown. All scale bars denote 500 μm. mDC = myeloid dendritic cells, Mono-mac = monocyte-derived macrophages, T reg = T regulatory cells.

Unsupervised clustering of snRNA-seq and multiome data identified immune, stromal, and tumour cell types (Fig 4b, Fig. S11e-f). The tumour cells were split into two subpopulations, denoted as tumour cluster 1 and tumour cluster 2, that segregated into spatially distinct compartments (Fig 4b-c, Fig. S11e). As copy number variation plays an important role in melanoma tumour evolution^46, 47^, we sought to identify if these transcriptional subpopulations represented distinct genetic clones. We inferred copy number alterations via inferCNV^48^, a standard scRNA-seq CNV inference tool, from the transcriptomes of each spatially mapped nucleus (Methods). Indeed, across both the snRNA-seq and the multiome data, we uncovered genomic differences consistent with the spatial and transcriptional separation between tumour cluster 1 and 2 (e.g. CNV on Chr6, Fig. 4d, Fig. S11g).

Our basic clustering analysis showed extensive T-cell infiltration into both tumour compartments (Fig. S11e, S12d); we wondered whether there might exist heterogeneous T-cell responses to these genetically distinct compartments. First, we enriched for TCR sequences in our 1,020 spatially positioned CD8 T-cell cDNA profiles, recovering 419 cells with alpha chains (279 unique), 761 cells with beta chains (410 unique), and 358 cells with paired alpha and beta chains (265 unique) (Table S7). We found a TCR beta clonotype that was significantly expanded in tumour compartment 2 compared to tumour compartment 1 (Fig. 4e, Fisher’s exact test, odds ratio = 6.8, *p* value = 1.1×10^-11^), in agreement with our previous report^49^. Given our high TCR pairing rate (Fig. S12f), we also noted tumour compartment 2 expansion of CD8 T cells with this beta chain and a paired alpha chain (Fisher’s exact test, odds ratio = 11.9, *p* value = 9.6×10^-6^). We observed CD8 T cells in tumour compartment 2 were upregulated in cytotoxic *GZMB* expression (Fig. S12g, Table S8). In addition to this T-cell variation, we noted decreased expression of MHC class I endogenous antigen presentation genes in tumour cluster 1 relative to tumour cluster 2 (Fig. S13, Gene set enrichment analysis GO:0002484, overlap ratio = 0.71, adj. *p* value = 6.6×10^-6^, Table S9), potentially contributing to differential T-cell clone infiltration between the tumour compartments. Thus, we observed a cytotoxic T-cell clone specifically infiltrating into a spatially and genetically distinct tumour compartment.

To further explore how chromatin accessibility and transcription informs tumour cell state and how this relates to the tumour microenvironment, we identified spatially-segregated differential gene expression and differential chromatin gene scores between tumour subpopulations (Fig. 4f, Table S10). *TNC* emerged as differentially expressed (log_2_FC = 2.1, adj. *p* value = 2.4×10^-61^, Methods) and differentially accessible by chromatin gene score (Wilcoxon Rank Sum test, log_2_FC = 0.81, adj. *p* value = 1.0×10^-12^) in tumour cluster 1 compared with tumour cluster 2 (Fig. 4g-h). We observed heterogeneity in *TNC* chromatin accessibility and gene expression within tumour cluster 1, which has previously been associated with a mesenchymal-like cell state^50–52^ therefore, we hypothesised tumour cluster 1 may comprise two cell states: melanocytic-like and mesenchymal-like. We scored tumour cells for melanocytic-like and mesenchymal-like cell states using genes previously implicated in this transition^50^. While tumour cluster 2 was largely a melanocytic-like population, we observed melanocytic-like and mesenchymal-like scores were negatively correlated and heterogeneous in tumour cluster 1 (Fig. 4i-j, Fig. S12h-i, Pearson’s *r* = -0.60, *p* value < 2.2×10^-16^). To uncover trans-acting factors associated with this transition, we first identified accessible transcription factor (TF) motifs that were correlated with mesenchymal-like score within tumour cluster 1 using ChromVar^53^ (Fig. 4k, x-axis, Table S11, adj. *p* value < 0.05); positively correlated TF motifs included FOS/JUN family members, which have previously been implicated in mesenchymal-like melanoma states, and IRF family TFs, while negatively correlated TF motifs included MITF, a factor involved in maintaining the melanocytic lineage^54, 55^. While such epigenomic signatures driving mesenchymal-like state have previously been identified in single cells, they have, to date, not been localised within tissues. To answer if such epigenetic signatures were spatially non-random, we performed spatial autocorrelation analysis on TF motif scores in the tumour cluster 1 compartment (Fig. 4k, y-axis, S12j). The top spatially autocorrelated TF motifs associated with mesenchymal-like state were JUN, FOS, and IRF family members with positive autocorrelation scores, suggesting these epigenomic signatures are locally clustered. Local clustering of epigenetic states is suggestive of inheritance of epigenetically reprogrammed states in cell division, or local signalling environmental drivers^42, 56, 57^.

## Discussion

Here we developed Slide-tags, a spatial single-nucleus genomics technology that is widely applicable to tissues spanning different scales, species, and disease states. We profiled Slide-tags nuclei isolated from mouse and human adult brain with snRNA-seq, showing indistinguishable RNA data quality and high spatial positioning accuracy, and discovering cell-type specific spatially varying genes across cortical layers. Applying Slide-tags snRNA-seq to densely-packed human tonsil enabled spatial contextualization of predicted receptor-ligand interactions. Finally, to demonstrate the multimodal capacity of Slide-tags, we simultaneously profiled the transcriptome, epigenome, and TCR repertoire of metastatic melanoma tissue, as well as inferred copy number variation from transcriptome data. We inferred copy number alterations from transcriptome data and revealed spatial immune cell differences between genomically distinct clones. In a cytogenetically homogenous subclone, we discovered two transitional tumour cell states and leveraged our single-nucleus spatial chromatin accessibility data to identify spatially auto-correlated transcription factor motifs likely to be participating in this mesenchymal-like transition.

Slide-tags offers several key advantages as a spatial genomics technology. First, it is easily imported into frozen tissue snRNA-seq experiments and allows the addition of spatially resolved data without requiring specialised equipment or sacrificing data quality. Second, the data is intrinsically single-cell resolution, without the need for deconvolution and segmentation, and is high sensitivity (2000-10000 UMIs/cell across our datasets). Third, the technology is high-throughput, enabling many tissue sections to be profiled at once, and coverage of larger tissue sections through the construction of bigger bead arrays. Fourth, Slide-tags is easily adapted to many different single-cell and single-nucleus methodologies. Beyond our demonstration of spatial snRNA-seq+snATAC-seq, we envision future adaptations of Slide-tags will enable the profiling of DNA^6, 58, 59^, additional epigenetic modifications^7, 60–63^, and proteins^64, 65^. Computational analyses of such data are uniquely enabled by the ability of Slide-tags to seamlessly leverage many existing single-cell computational workflows (e.g. Seurat^23^, InferCNV^48, 66^, ArchR^66^).

While immediately useful in many applications, Slide-tags could be improved in two key ways. First, our method only assays a subset of nuclei in a tissue section. We estimate that the combination of dissociation and microfluidic losses during nuclei barcoding collectively account for ∼75% of the nuclei lost. This represents an avenue for significant improvement through tissue-specific optimizations to the dissociation, and improved droplet microfluidics or, potentially, microfluidic free single-nucleus methods that may barcode nuclei more efficiently^67^. Second, Slide-tags is currently limited to single nucleus sequencing methods, primarily due to the ease of recovering nuclei from frozen tissues. Some methodologies strongly benefit from single-cell data, such as lineage tracing using mitochondrial genomic variants^68, 69^, and quantification of transcriptional kinetics^70^. Future iterations of our technology may be compatible with tagging whole single cells. Nonetheless, for routine tissue profiling, our current default approach is snRNA-seq (versus scRNA-seq), owing to advantages in protocol flexibility, increased nuclei yields, reduced tissue dissociation artefacts, and improvements to cell sampling bias^71^.

In recent years, a common experimental paradigm has evolved which pairs the collection of single-cell (or single-nucleus) data with spatial data to discover cell types, compare across conditions, and discover spatial patterns within and across these types. Slide-tags represents a method to merge these experimental modalities into a unified approach, integrating the ascertainment of cytoarchitectural features with the standard collection of single-cell sequencing data. By importing the single-cell sequencing toolkit into the spatial repertoire, Slide-tags will serve as an invaluable tool to study tissue biology across organisms, ages, and diseases.

## Acknowledgments

We thank Nina Sachdev, Jonah Langlieb, Gordon Fishell, Sherry Wu, Nir Hacohen, Arnav Mehta, the Macosko and Chen laboratories, for helpful discussions. We thank David Davis and William Scott (University of Miami Brain Endowment Bank), for their contribution of human postmortem brain tissue. Components of our figures were created with BioRender.com. We thank the patients and their families for their invaluable donations to science, making this work possible. This work was supported by the National Institutes of Health (grant nos. R01HG010647 and UH3CA246632 to E.Z.M. and F.C.). F.C. also acknowledges support from the Searle Scholars Award, the Burroughs Wellcome Fund CASI award, and the Merkin Institute. A.J.C.R is supported by an EMBO Postdoctoral Fellowship.

## Author contributions

A.J.C.R., J.A.W., N.M.N., E.Z.M., and F.C. conceived the study. A.J.C.R., J.A.W., and N.M.N. developed the Slide-tags protocol and performed experiments with help from G.M., K.B., C.V., and V.S. L.T. wrote barcode matching code. A.J.C.R., J.A.W., and M.S. performed analyses with help from S.K and F.C. V.K. synthesised the barcoded beads. C.W. and C.Y. provided the melanoma biopsy tissue. A.J.C.R., J.A.W., E.Z.M., and F.C. wrote the paper with contributions from all authors.

## Data and code availability

Slide-tags datasets have been deposited on the Broad Institute Single Cell Portal, under the following accession numbers: <u>mouse brain</u> (SCP2162), <u>mouse embryonic brain</u> (SCP2170), <u>human brain</u> (SCP2167), <u>human tonsil</u> (SCP2169), <u>human melanoma</u> (SCP2171), <u>human</u> <u>melanoma multiome</u> (SCP2176). Mouse raw sequencing data will be available from the Sequence Read Archive. Raw and processed data to support the findings of this study will be deposited in GEO. Code for processing spatial sequencing libraries is available on Github: https://github.com/broadchenf/Slide-tags.

## Materials and Methods

### 1 Experimental methods

#### 1.1 Sample information and processing

##### Mouse brain

###### Animal housing

Animals were group-housed with a 12-hour light-dark schedule and allowed to acclimate to their housing environment for two weeks post arrival. All procedures involving animals at the Broad Institute were conducted in accordance with the US National Institutes of Health Guide for the Care and Use of Laboratory Animals under protocol number 0120-09-16 and approved by the Broad Institutional Animal Care and Use Committee.

###### Brain preparation

At 56 days of age, C57BL/6J mice were anaesthetised by administration of isoflurane in a gas chamber flowing 3% isoflurane for 1 minute. Anaesthesia was confirmed by checking for a negative tail pinch response. Animals were moved to a dissection tray and anaesthesia was prolonged via a nose cone flowing 3% isoflurane for the duration of the procedure. Transcardial perfusions were performed with ice cold pH 7.4 HEPES buffer containing 110 mM NaCl, 10 mM HEPES, 25 mM glucose, 75 mM sucrose, 7.5 mM MgCl_2_, and 2.5 mM KCl to remove blood from brain and other organs sampled. For use in regional tissue dissections, the brain was removed immediately and frozen for 3 minutes in liquid nitrogen vapour and then moved to -80 °C for long term storage.

Whole C57 mouse embryos at E14 (MF-104-14-Ser) were purchased from Zyagen and stored at −80 °C until use. A pregnant mouse was perfused with PBS prior to harvesting and snap freezing of the whole embryo.

###### Human brain

Postmortem autopsy tissue (Brodmann area 9 cortex) from a healthy, aged, female, control case was obtained from the University of Miami Brain Endowment Bank at the Miller School of Medicine. Tissue was collected in accordance with the standard patient informed consent procedures of the Brain Endowment Bank in effect at the time of collection and subject to approval or an exemption determination by their Institutional Review Board. Use of the tissue at the Broad Institute was approved by the Office of Research Subject Protection project NHSR-4235. This cortical specimen was stored at -80 °C until use following equilibration at -20 °C in the cryostat. As a quality control step, tissue architecture was assessed by Nissl staining following frozen sectioning at 20 µm, and RNA integrity was determined using trizol extraction followed by RIN assay via the Agilent RNA nano 6000 bioanalyzer method (RIN = 7.2).

###### Human tonsil

Anonymized excess tissue specimens were obtained from a patient who underwent a palatine tonsillectomy procedure for tonsillar enlargement. The specimens were embedded in OCT, snap-frozen and stored at -80°C. As a quality control step, tissue architecture was assessed by hematoxylin and eosin staining, and RNA integrity was determined using the Tapestation RNA ScreenTape system (RIN^e^ > 7.5). Use of the tissue at the Broad Institute was approved by the Office of Research Subject Protection project IRB-6429.

###### Human metastatic melanoma

Specimens were acquired from a patient who underwent axillary lymphadenectomy for metastatic BRAF-mutant melanoma prior to starting PD-1 inhibitor. The specimen was embedded in OCT, snap frozen following surgery, and stored at -80 °C. Use of the tissue at the Broad Institute was approved by the Office of Research Subject Protection project NHSR-4182.

#### 1.2 Histological processing

For sections that were stained using Nissl, glass-mounted frozen tissue sections (10 or 20 µm) were equilibrated to RT and excess condensate was wiped off. Sections were fixed in 70% ethanol for 2 min, followed by rehydration in ultrapure water for 30 s. Excess water was wiped off and slides were stained with Arcturus Histogene Solution (ThermoFisher, no. 12241-05) for 4 min. Excess dye was tapped off and slides were rehydrated in water for 10 s for destaining. Slides were sequentially fixed in 70, 90 and 100 % ethanol for 30 s, 10 s and 1 min, respectively, post-fixed in xylene solution for 1 min then mounted with Fisher Chemical Permount (no. SP15-100) and coverslipped. Images were acquired with a Keyence BZ-800XE microscope under a Nikon Apo 10x objective or the the Leica Aperio VERSA Brightfield, Fluorescence & FISH Digital Pathology Scanner under a 10x objective.

For sections that were stained using hematoxylin and eosin H&E, glass-mounted frozen tissue sections (10 or 20 µm) were equilibrated to RT and excess condensate was wiped off. Sections were dipped in xylene, processed through a graded ethanol series, and stained with hematoxylin. The nuclei were “blued” by treatment with a weakly alkaline solution, and washed with water. Sections were stained with eosin, processed through a graded ethanol series, xylene, dehydrated, and coverslipped. Brightfield images were taken using the Leica Aperio VERSA Brightfield, Fluorescence & FISH Digital Pathology Scanner under a 10x objective.

#### 1.3 Barcoded bead synthesis, array fabrication, and sequencing

PLRP-S resin (1000 A, 10-μm; Agilent Technologies, PL1412-4102) was used for the barcoded oligonucleotide synthesis. The loading of the non-cleavable linker on resin was adjusted to approximately 30 µmol/g. The Akta OligoPilot 10 oligonucleotide synthesizer was used for synthesis (850 mg scale). The PC linker (cat. no. 10-4920-90) and reverse phosphoramidites (10-0001, 10-9201, 10-0301, and 10-5101-10) were purchased from Glen Research. A 0.1 M solution of phosphoramidites was prepared in anhydrous acetonitrile (ACN) and 0.3 M BMT (BI0166-1005, Sigma-Aldrich) was used as an activator for coupling (single coupling, 6 min). Two capping steps (before and after oxidation) were performed with Cap A (BI0224-0505, Sigma-Aldrich) and Cap B (B1:B2 1:1; BI0347-0505, BI0349-0505 Sigma-Aldrich) reagents. For the 6.3 mL column, capping was performed by 1 CV or 1.5 CV with one min and for 1.2 mL column, 2 CV for 0.5 min. The oxidation (5 equiv) was carried out with 0.05 M iodine in pyridine (BI0424-1005, Sigma-Aldrich). The detritylation step was performed using 3% dichloroacetic acid in toluene (BI0832-2505, Sigma-Aldrich).

After the oligonucleotide synthesis, the protecting groups were removed by incubating the resin in 40% aqueous methylamine for 24 hr at room temperature (20 mg resin/ 2mL). The beads were washed twice with water (1 mL), three times with methanol (1 mL), three times with 1:1 acetonitrile: water, and three times with acetonitrile (1 mL). Finally, beads were washed three times with 10 mM Tris buffer pH 7.5 containing 0.01% tween-20 and stored in the same buffer at 4 °C. It was observed that oligos were released in the buffer if the beads were stored for long periods of time. In order to remove the released oligos, beads were washed with 70% acetonitrile/ water and resuspended in storage buffer.

Synthesised sequences for the Slide-tags experiments (PC in the sequences denote photocleavable linker):

1) Incorporation of capture sequence by ligation: blue colour letters denote the region that is complementary to the sequence of the 10x Gel beads.

5’-TTT_PC_GCCGGTAATACGACTCACTATAGGGCTACACGACGCTCTTCCGATCTJJJJJJJJTCTT CAGCGTTCCCGAGAJJJJJJJNNNNNNNVVGCTCGGACACATGGGCG-3’

10X FB1 extension: 5’-GAGCTTTGCTAACGGTCGAGGCTTTAAGGCCGGTCCTAGCAA-3’ Splint: 3’-CTGTGTACCCGCCTCGAAACGATTGC-5’

2) Direct synthesis of capture sequence on beads:

5’-TTT-PC-GTGACTGGAGTTCAGACGTGTGCTCTTCCGATCTJJJJJJJJTCTTCAGCGTTCCCGA GAJJJJJJJNNNNNNNVVGCTTTAAGGCCGGTCCTAGCAA-3’

3) Poly A beads:

5’-TTT-PC-GTGACTGGAGTTCAGACGTGTGCTCTTCCGATCTJJJJJJJJTCTTCAGCGTTCCCGA GAJJJJJJJNNNNNNNVVA30

Array preparation and sequencing were performed as described previously^22^.

#### 1.4 Slide-tags procedure

Fresh frozen tissues were cryo-sectioned to 20 μm on a cryostat (CM1950, Leica) at -16 °C. Pre-cooled 2 mm circular (3331P/25, Integra), 3 mm circular (3332P/25, Integra), or 5.5 mm square custom-made biopsy punches were used to isolate regions of interest from tissue sections. The punched tissue regions were then placed on the puck, ensuring there were no folds. A finger was placed on the bottom of the puck to melt the tissue whilst trying to prevent rolling. Immediately this puck was placed on the glass slide and placed on ice, and 6-10 µL of dissociation buffer (82 mM Na_2_SO_4_, 30 mM K_2_SO_4_, 10 mM Glucose, 10mM HEPES, 5 mM MgCl_2_) was placed on top of the puck so that the buffer covered the whole puck. The puck was then placed under a UV (365 nm) light source (0.42 mW/mm^2^, Thorlabs, M365LP1-C5, Thorlabs, LEDD1B) for 30 s (TAGS beads), or 3 mins (SLAC beads), in order to cleave the same amount of spatial barcode oligonucleotides between bead designs (Fig. S2h). After photo-cleavage, the puck was incubated for 7.5 mins (TAGS beads) or 5 mins (SLAC beads) and then placed into a 12-well plate (Corning, 3512). Using a 200 µL pipette, 10 x 200 µL aliquots of extraction buffer (Dissociation Buffer, 1% Kollidon VA64, 1% Triton X100, 0.01% BSA, 666 units/mL RNase-inhibitor (Biosearch technologies, 30281-1)) were dispensed onto the puck for a total volume of 2 mL. Dispensed extraction buffer was triturated up and down on the puck for 10-15 times to release the tissue. This step was repeated until the tissue was completely removed from the puck. The puck was removed, and mechanical dissociation of the supernatant was performed using 1 mL pipette 20-25 times trituration to fully dissociate the tissue. Dissociated nuclei were removed from the well and the well was rinsed twice with 1 mL of wash buffer (82 mM Na_2_SO_4_, 30 mM K_2_SO_4_, 10 mM Glucose, 10mM HEPES, 5 mM MgCl_2_, 50 µl of RNase-inhibitor (Biosearch technologies, 30281-1)) which was added to nuclei suspension. Wash buffer was added to the tube to a final volume of 20 mL. This 20 mL was mixed and divided equally into another 50 mL falcon tube. Nuclei were spun in a pre-cooled swinging bucket centrifuge at 600 g for 10 min at 4 °C. After centrifugation, 19.5 mL of supernatant was removed, leaving 500 µL in each tube. The pellet was resuspended and pooled. This pooled suspension was then filtered using a pre-cooled 40 µm cell strainer (Corning, 431750). DAPI (Thermo Fisher Scientific, 62248) was added to the filtered solution at a 1:1000 dilution and incubated for 5-7 mins at 4 °C. This was then centrifuged at 200 g for 10 mins at 4 °C. The supernatant was removed, leaving 50 µL of pellet. The pellet was resuspended and nuclei were counted manually using a C-Chip Fuchs-Rosenthal disposable hemocytometer (INCYTO, DHC-F01-5).

#### 1.5 Sequencing library preparation

##### snRNA-seq library preparation

For Slide-tags snRNA-seq experiments, 43.3 µL of counted nuclei were loaded into the 10x Genomics Chromium controller using the Chromium Next GEM Single Cell 3’ Kit v3.1 (10x Genomics, PN-1000268). The Chromium Next GEM Single Cell 3’ Reagent Kits v3.1 (Dual Index) with Feature Barcode technology for Cell Surface Protein CG000317 was used according to the manufacturer’s recommendations with slight modifications. Spatial barcode libraries were prepared as Cell Surface Protein Library preparations. The number of PCR cycles used for the index PCR step in the Cell Surface Protein Library preparation (step 4.1f) for 5.5×5.5 mm TAGS arrays was 7; for 3 mm diameter TAGS arrays the number of cycles was 9.

For the mouse brain sample, ligated pucks (see sequence in section 1.3) were used for spatial barcoding. For this sample, a custom PCR protocol was used instead of step 4.1: 10 uL of cleaned supernatant from step 2.3, 50 µL NEBNext High-Fidelity 2X PCR Master Mix (NEB, M0541S), 2.5 µL STAG_P701_NEX (10 uM), 2.5 µL 10 μM P5-Truseq Hybrid oligo, 35 µL UltraPure DNase/RNase-Free Distilled Water (Invitrogen, 10977015). In this sample, 10 PCR cycles were performed according to the manufacturer’s recommendations.

##### snATAC-seq and snRNA-seq library preparation

For Slide-tags multiomic snATAC-seq and snRNA-seq experiments, 43.3 µL of counted nuclei were loaded into the 10x Genomics Chromium controller using the Chromium Next GEM Single Cell Multiome ATAC + Gene Expression Reagent Bundle (10x Genomics, PN-1000283). The Chromium Next GEM Single Cell Multiome ATAC + Gene Expression CG000338 Rev F user guide was used according to the manufacturer’s recommendations with slight modifications. During step 4.1, 1 uL of 0.329 uM spike-in primer (5’-GTGACTGGAGTTCAGACGT-3’) was added. For spatial barcode libraries, a custom PCR protocol was used: 5 uL of cleaned supernatant from step 4.3, 50 µL NEBNext High-Fidelity 2X PCR Master Mix (NEB, M0541S), 2.5 µL 10 μM STAG_iP7_a1 oligo (5’-CAAGCAGAAGACGGCATACGAGATATTTACCGCAGTGACTGGAGTTCAGACGT*G*T-3’), 2.5 µL 10 μM P5-STAG_ip5_a1 oligo (5’-AATGATACGGCGACCACCGAGATCTACACGACAATAAAGACACTCTTTCCCTACACGACGC* T*C-3’), 40 µL UltraPure DNase/RNase-Free Distilled Water (Invitrogen, 10977015). In this sample, 15 PCR cycles were performed according to the protocol used in The Chromium Next GEM Single Cell 3’ Reagent Kits v3.1 (Dual Index) with Feature Barcode technology for Cell Surface Protein CG000317 Rev C user guide step 4.1.

##### T cell receptor enrichment and library preparation

We enriched TCRs from Slide-tags multiome cDNA as previously described^49^ with the following modifications (https://www.protocols.io/view/slide-tcr-seq-v3-ivt-n92ldp6w8l5b/v2).

#### 1.6 Sequencing

We sequenced scRNA-seq and spatial barcode libraries on an Illumina Nextseq 1000 instrument using a p2 100 cycle kit (Illumina, 20046811). For some libraries, resequencing was performed to improve sequencing depth, on an Illumina Novaseq instrument using the S Prime platform.

### 2 Slide-tags data preprocessing

#### 2.1 snRNA-seq data

We used Cell Ranger v6.1.2^2^ mkfastq (10x Genomics) to generate demultiplexed FASTQ files from the raw sequencing reads. We aligned these reads to either the human GRCh38 or mouse mm10 genome whilst including intronic reads with --include-introns, and quantified gene counts as UMIs using Cell Ranger count (10x Genomics). For mouse embryo, human brain, tonsil, and melanoma, we used CellBender v0.2.0 for background noise correction and cell calling^72^, setting --expected-cells to the number of Cell Ranger cell calls, --total-droplets-included to 40,000, and --learning-rate to 0.00005 (only when default parameters were insufficient to produce cell probabilities calls of majority zero and one).

#### 2.2 Multiomic snATAC-seq and snRNA-seq data

We used Cell Ranger-arc v2.0.2 mkfastq (10x Genomics) to generate demultiplexed FASTQ files from the raw sequencing reads. We aligned these reads to the human GRCh38 genome, and quantified gene counts as UMIs using Cell Ranger-arc count (10x Genomics). For the gene expression data, we then used CellBender for background noise correction and cell calling as described above.

#### 2.3 Spatial barcode data

After creating demultiplexed FASTQ files, we grepped for reads containing the spatial barcode universal primer constant sequence. We then downsampled the spatial barcode-containing FASTQ file to 25 million reads using seqtk v1.3-r106 for computational efficiency and consistency across runs. We then matched candidate cell barcodes in the spatial barcode FASTQ file with true cell barcodes outputted from either Cell Ranger v6.1.2 or CellBender^72^ (Table S12), generating a data frame of candidate spatial barcode sequences per true cell barcode. From this data frame, we matched candidate spatial barcode sequences with a whitelist of *in situ* sequenced spatial barcodes, assigning each true spatial barcode a spatial coordinate.

#### 2.4 Assignment of spatial locations to nuclei

The set of spatial barcodes per cell barcode can be used for spatial positioning of nuclei. To do so, we first read the processed barcodes into R and removed spatial barcodes with nUMI > 256 as these likely represented beads that have been dislodged from the glass slide and encapsulated in droplets with nuclei (data not shown). We then used nUMI-weighted density-based spatial clustering of applications with noise (DBSCAN)^73, 74^ v1.1-11 to distinguish “signal” spatial barcodes (those likely to provide value in positioning nuclei) from background “noise” spatial barcodes (those likely to confound nuclei positioning). DBSCAN outputs a cluster assignment for each spatial barcode. Cluster = 0 corresponds to “noise” spatial barcodes without a clear spatial distribution, and cluster numbers above zero correspond to “signal” spatial barcodes with discrete spatial distributions. We did not assign spatial positions to nuclei with all spatial barcodes denoted noise, or to nuclei with multiple signal clusters. From the remaining nuclei with one distinct spatial barcode signal cluster, we took an nUMI-weighted centroid of spatial barcode coordinates in the signal cluster. DBSCAN required two parameters as input: *minPts* and *eps* (i.e., radius). To determine the optimal parameter set for each Slide-tags run, we iterated through *minPts* parameters from *minPts* = 3 to *minPts* = 15 under a constant *eps* = 50 and chose the parameter set with the highest proportion of nuclei that are assigned a spatial position (one DBSCAN signal cluster).

#### 2.5 T cell receptor sequences

TCR sequences were identified using MiXCR v4.1.0^75, 76^ and assigned to cell barcodes using a hamming distance 1 collapse.

### 3 Mouse brain analysis

#### 3.1 Quality control and cell type assignment

The output generated by Cell Ranger was read into R (4.1.1) using Seurat (4.3.0)^23^. We normalised the total UMIs per nucleus to 10,000 (CP10K) and log-transformed these values to report gene expression as E = log(CP10K + 1). We identified the top 2000 highly variable genes after using variance-stabilizing transformation correction^77^. All gene expression values were scaled and centred. For visualisation in two dimensions, we embedded nuclei in a Uniform Manifold Approximation and Projection (UMAP)^78^ using the top 30 PCs, with: number of neighbours = 40, min_dist = 0.3, spread = 15, local connectivity = 12, and the cosine distance metric. We identified shared nearest neighbours using the top 30 principal components. Clusters of similar cells were detected using the Louvain method for community detection, implemented using *FindClusters*, with a resolution = 0.8. Each cell was then assigned a predicted identity based on mapping to a mouse adult brain reference dataset^18^, using FindTransferAnchors and then TransferData, with the first 25 PCs in both cases. For each computed cell cluster, an identity was assigned using the highest proportion of transferred labels, and confirmed using known markers genes

#### 3.2 Assessment of spatial positioning accuracy

##### Spatial barcode metrics calculations

We measured the accuracy of spatial positioning for the 839 cell barcodes corresponding to high-quality mapped cells in our mouse hippocampus dataset (Figure 1). For each of these cells, we used the spatial barcodes belonging to the DBSCAN singlet cluster and calculated the standard error for both x and y coordinates using:

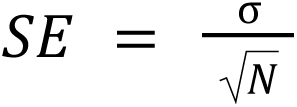

Where: *N* is the number of spatial barcode UMIs in the cluster, and σ is the standard deviation of each of the spatial barcode UMIs from the centroid of the cluster.

In addition to the SE, other metrics were calculated for each DBSCAN singlet cluster. Namely, the geometric mean distance of spatial barcodes from the centroid:

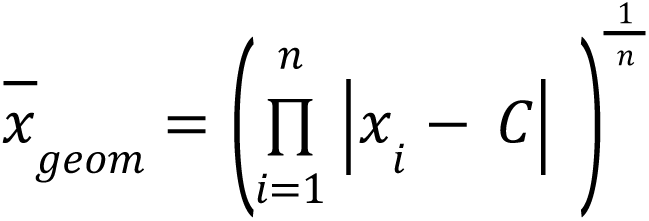

Where: *n* is the number of spatial barcode UMIs in the cluster, and |*x_i_* − *C*| is the absolute distance between each spatial barcode UMI and the cluster centroid.

For each cell that had only a single DBSCAN cluster, additional metrics were calculated (Fig. S2d). The total number of unique spatial barcode sequences, and spatial barcode UMIs associated with each cell was calculated, regardless of whether it was in the singlet DBSCAN cluster or not. The ratio of spatial barcode UMIs within and outside the DBSCAN singlet cluster was then calculated as the proportion of signal spatial barcodes per cell.

##### CA1 width analysis

A serial section of the profiled region was stained using Nissl and imaged. Cells were segmented from this image via watershed segmentation in Matlab (Release 2021b) and the centroid of each segment was calculated. Next, these coordinates were read into R and DBSCAN was used to isolate cells belonging to the CA1 region, with the following parameters: eps = 35, minPts = 20. The image region was cropped to match that of the profiled Slide-tags region. For both datasets, a 10th (Nissl) or 9th (Slide-tags) order linear model was fitted through these points, generating a central curve. For each spatial barcode UMI, the nearest neighbour on this curve in euclidean space was determined and the distance from these two points was recorded as the distance from the fitted line.

#### 3.3 Comparison of Slide-tags snRNA-seq vs. snRNA-seq data

For each sample, cellranger was run as above, and the outputs were run through cellranger aggr (v6.1.2), in order to account for differences in sequencing depth per cell. The result was a combined matrix of 25,158 nuclei, with 25,107 mean reads per cell, 2,309 median UMIs per cell, and 1,438 median genes per cell. The filtered feature-barcode matrix generated by Cell Ranger was then read into R (4.1.1) using Seurat (4.3.0)^23^. We normalised the total UMIs per nucleus to 10,000 (CP10K) and log-transformed these values to report gene expression as E = log(CP10K + 1). We identified the top 2000 highly variable genes after using variance-stabilizing transformation correction^77^. All gene expression values were scaled and centred. For visualisation in two dimensions, we embedded nuclei in a Uniform Manifold Approximation and Projection (UMAP)^78^ using the top 40 PCs, with: number of neighbours = 40, min_dist = 0.3, spread = 15, local connectivity = 12, and the cosine distance metric. We identified shared nearest neighbours using the top 40 principal components. Clusters of similar cells were detected using the Louvain method for community detection, implemented using *FindClusters*, with a resolution = 1. Each cell was then assigned a predicted identity based on mapping to a mouse adult brain reference dataset^18^, using FindTransferAnchors and then TransferData, with the first 25 PCs in both cases. These cell type designations were then used for comparative analysis going forward. Cells designated “Unk_1” or “Unk_2” were removed from the analysis as these cells showed low quality metrics and were not interpretable labels.

### 4 Mouse embryonic brain at E14 analysis

The output generated by Cell Ranger was read into R (4.1.1) using Seurat (4.3.0)^23^. We normalised the total UMIs per nucleus to 10,000 (CP10K) and log-transformed these values to report gene expression as E = log(CP10K + 1). We identified the top 2000 highly variable genes after using variance-stabilizing transformation correction^77^. All gene expression values were scaled and centred. For visualisation in two dimensions, we embedded nuclei in a Uniform Manifold Approximation and Projection (UMAP)^78^ using the top 30 PCs, with: number of neighbours = 40, min_dist = 0.3, spread = 15, local connectivity = 12, and the cosine distance metric. We identified shared nearest neighbours using the top 30 principal components. Clusters of similar cells were detected using the Louvain method for community detection, implemented using *FindClusters*, with a resolution = 0.8. Each cell was then assigned a predicted identity based on mapping to a mouse embryo at E14 reference dataset^20^, using FindTransferAnchors and then TransferData, with the first 25 PCs in both cases. For each computed cell cluster, an identity was assigned using the highest proportion of transferred labels, and confirmed using known marker genes.

### 5 Human brain analysis

#### 5.1 Quality control and cell type assignment

The output generated by Cell Ranger was filtered by CellBender and read into R (4.2.2). The matrix was subsetted down to cells that had exactly one DBSCAN location, which were then loaded into Seurat (4.3.0)^23^ to perform normalisation, finding variable features, scaling, PCA, finding neighbours (dims=30), finding clusters, and creating a UMAP, all with default parameters (unless specified otherwise). Each cluster was assigned a cell class (Excitatory neuron, Inhibitory neuron, Oligodendrocyte (Oligo), Oligo precursor cell (OPC), Astrocyte, Endothelial cell, Microglia) by plotting canonical cell type marker genes on the UMAP and manually assigning each cluster a cell type. Clusters were removed if the average percentage of mitochondrial reads exceeded 5% or if the top 10 differentially expressed genes between the cluster and other clusters of the same cell type belonged to a different cell class. Subsequently, excitatory and inhibitory neuron subtypes were mapped from a published human cortex dataset^28^ by label transfer using Harmony v0.1.1 and spatially plotted in Fig. S6.

#### 5.2 Identification of layers and layer-dependent gene expression

The layer assignment of each cell (L1-2, L3-5, L6, WM) was calculated by manually drawing boundaries between the layer-specific mapped neuron subtypes and assigning each cell a label depending on which two boundaries it was between. The numerical laminar coordinate was then calculated by taking the Euclidean distance of each cell to the nearest boundary and dividing it by the sum of the distances to the two neighbouring boundaries, adding a constant factor depending on the layer assignment. Each gene was assigned a spatial variation score by computing the kernelized density of the gene expression along the laminar coordinate using a uniform kernel and taking the difference between the highest and lowest expression density values (Table S1). Complex gradients were found by taking the intersection of each cell type’s spatially variable gene list, and a visually-selected interesting subset is shown in Fig. 2l.

Gene ontology analysis was performed on all genes with a spatial variation score above 0.5 using EnrichGO from clusterProfiler 4.6.0 (default parameters) and using annotations from org.Hs.eg.db v3.16.0 (Table S2). For display in Fig. S9, the full list was subsetted to show all terms with an adjusted p-value <0.05 in more than one cell type and terms that did not have ancestor GO terms with p-value <0.05 in the same cell types. For display in Fig. 2k, the terms were further subsetted to only include terms in the Biological Process (BP) ontology with an adjusted p-value below 0.0001 in at least one cell type.

Genes with a spatial variation score above 0.8, or above 0.55 with a minimum expression below 1, were shown in the heatmaps in Fig. 2i-j, S5d-e. Genes with a spatial variation score above 0.8, or above 0.60 with a minimum expression below 1 were spatially plotted in Fig. S7-8.

### 6 Tonsil analysis

### 6.1 Quality control and cell type assignment

The output generated by Cell Ranger and filtered by CellBender was read into R (4.1.1) using Seurat (4.3.0)^23^. We normalised the total UMIs per nucleus to 10,000 (CP10K) and log-transformed these values to report gene expression as E = log(CP10K + 1). We identified the top 2000 highly variable genes after using variance-stabilising transformation correction^77^. All gene expression values were scaled and centred. For visualisation in two dimensions, we embedded nuclei in a Uniform Manifold Approximation and Projection (UMAP)^78^ using the top 30 PCs, with: number of neighbours =30, min_dist = 0.3, spread =1, local connectivity = 1, and the cosine distance metric. We identified shared nearest neighbours using the top 30 principal components. Clusters of similar cells were detected using the Louvain method for community detection, implemented using *FindClusters*, with a resolution = 1. Annotation of *de novo* clusters was aided by marker genes and Azumith^23^ reference-based mapping from the human tonsil atlas^79^.

### 6.2 Spatially varying gene expression

Significantly nonrandom genes were discovered in germinal centre B cells as described previously^10^. Briefly, for each single-nucleus assigned as a germinal centre B cell that was positioned in one of the four largest germinal centres we profiled, we first calculated the matrix of pairwise Euclidean distances between cells for each germinal centre individually. We then compared the distribution of pairwise distances between the cells expressing at least one count of that transcript to the distribution of pairwise distances between an identical number of cells, sampled randomly from all mapped beads within the set with probability proportional to the total number of UMIs per cell. Specifically, we generated 1000 such random samples, and for each sample calculated the distribution of pairwise distances. We then calculated the average distribution of pairwise distances, averaged across all 1000 samples. Finally, we calculated the L1 norm between the distribution of pairwise distances for the true sample of cells and the average distribution. We defined p to be the fraction of random samples having distributions closer to the average distribution (under the L1 norm) than the true sample. We calculated an Z-score for the true sample given the distribution distances from the average distribution of random samples. Finally, we aggregated *p* values for spatial variation from each of the four tested germinal centres using Fisher’s method.

We intersected our computed spatially varying genes with genes previously implicated in germinal centre zone distinction^80^. We calculated percent variance in gene expression space and plotted against spatial effect size from our spatial permutation test to identify genes with relatively low gene expression variance but high spatial variance.

### 6.3 Germinal centre zonation

We used spatially varying genes (*p* value < 0.05) identified as described above to classify germinal centre B cells into light zone, dark zone, and transitional states. Specifically, we subsetted our data to germinal centre B cells, re-scaled and re-centred values, and ran PCA on the 1068 significant spatially varying genes. We then identified shared nearest neighbours using the top 15 principal components. Clusters of similar cells were detected using the Louvain method for community detection, implemented using *FindClusters*, with resolution = 0.4. We annotated clusters as light zone, dark zone, and transitional states using marker genes and Azumith^23^ reference-based mapping from the human tonsil atlas^79^.

After classifying germinal centre B cells into states, we spatially segmented germinal centres into light zones and dark zones using dark zone B cell spatial density. We ran DBSCAN^73^ on dark zone B cells of the two largest germinal centres, using *eps* = 60 and *minPts* = 6 for the largest germinal centre, and *eps* = 60 and *minPts* = 10 for the second largest germinal centre. We considered cells within the top DBSCAN cluster to constitute the dark zone and segmented around the outer cells. The remaining cells in both germinal centres were considered to be in the light zone and segmentation borders were drawn accordingly. We tested for zone bias of T follicular helper cells and follicular dendritic cells using *chisq.test* from the stats package in R (4.2.2).

### 6.4 Spatial receptor-ligand prediction

To detect receptor-ligand interactions between cell-type pairs, we computed a receptor-ligand score based on a spatial correlation index^81^, SCI, which we defined as:

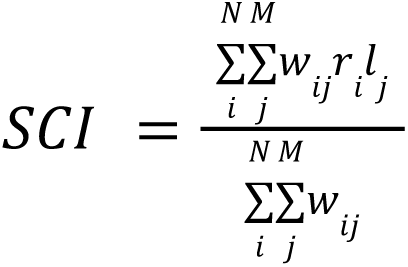

between N cells of “sender cell type” expressing receptor r and M cells of “receiver cell type” expressing ligand l, where expression is sctransform counts^82^. We defined the spatial weights matrix of dimensionality NxM as an adjacency matrix, denoting 1 for when sender cell i is within 100 um of receiver cell j and 0 otherwise. We first ran LIANA^37^ (0.1.12) to generate a putative list of receptor-ligand interactions between cell-type pairs in a spatial agnostic way, filtering to receptor-ligand interactions that are expressed in at least 50 cells of sender and receiver cell types (log CPM > 0), or in 30% of sender and receiver cells. We then computed a spatial correlation index for each receptor-ligand interaction to determine if the receptor and ligand are spatially co-expressed in a given cell-type pair.

To determine the spatial significance of a receptor-ligand score, we employed an adaptive spatial permutation test, running 1000 permutations for each receptor-ligand interaction. In each permutation, we randomly permuted the spatial locations of cells within a given cell-type. For interactions that have a nominal *p* value less than or equal to 0.005, we ran an additional 9000 permutations. We corrected for multiple hypothesis testing using the Benjamini-Hochberg procedure. We also computed the log-fold change between the observed SCI statistic and the median SCI statistic of the empirical null distribution. This allowed us to compare SCI log-fold change values between receptor-ligand interactions for different cell types without explicitly correcting for the number of cells of each cell type.

### 6.5 Spatial contextualization of receptor-ligand interactions

To spatially contextualise receptor-ligand interactions, we decomposed spatial correlation indices for each significant interaction between germinal centre B cells, T follicular helper cells, and follicular dendritic cells (adj. *p* value < 0.05) into interaction intensity scores for individual cells^83^. These decomposed scores reflect each individual cell’s contribution to the total spatial correlation index, defined as follows for receiving cell i and vice-vera’s for sender cell j:

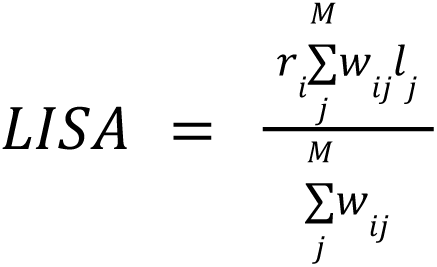

We tested germinal centre zone specificity via *wilcox.test* in R comparing interaction intensity scores of the receptor of each cell between dark zones and light zones. We corrected *p* values using the Benjamini-Hochberg method. Zone-specific receptor expression was tested using SCTransformed expression values compared between dark zones and light zones also using *wilcox.test* in R.

### 7 Melanoma analysis

#### 7.1 Quality control and cell type assignment

##### snRNA-seq data

The Cell Ranger output was filtered by CellBender and read into R (4.1.1) using Seurat (4.3.0)^23^. We normalised the total UMIs per nucleus to 10,000 (CP10K) and log-transformed these values to report gene expression as E = log(CP10K + 1). We identified the top 2000 highly variable genes after using variance-stabilising transformation correction^77^. All gene expression values were scaled and centred. For visualisation in two dimensions, we embedded nuclei in a Uniform Manifold Approximation and Projection (UMAP)^78^ using the top 30 PCs, with: number of neighbours =30, min_dist = 0.3, spread =1, local connectivity = 1, and the cosine distance metric. We identified shared nearest neighbours using the top 30 principal components. Clusters of similar cells were detected using the Louvain method for community detection, implemented using *FindClusters*, with a resolution = 1. Annotation of *de novo* clusters was aided by marker genes.

##### Multiome ATAC & snRNA-seq data

The RNA expression matrix generated by Cell Ranger was read into R (4.1.1) using Seurat^23^. The ATAC filtered feature-barcode matrix generated by Cell Ranger was read into R (4.1.1) using Signac (1.9.0)^84^, and added as its own assay slot in the Seurat object containing RNA expression counts. Peaks were recalled using the CallPeaks function, which uses MACS2 (2.2.7.1)^85^, across all cells. Fragments were mapped to the MACS2-called peaks and assigned to nuclei using the FeatureMatrix function in Signac. Peaks in non-standard chromosomes were removed using keepStandardChromosomes from GenomeInfoDb (1.35.15)^86^, and problematic regions of the hg38 genome were removed using subsetByOverlaps according to the blacklist available at: https://github.com/Boyle-Lab/Blacklist87. This final peaks-barcode matrix was then added to the “peaks” assay within the Seurat object.

For cell type annotation, the snRNA-seq data from the multiome experiment was normalised for the total UMIs per nucleus to 10,000 (CP10K) and log-transformed to report gene expression as E = log(CP10K + 1). The top 2000 highly variable genes were identified after using variance-stabilising transformation correction^77^. We then integrated the gene expression data from Slide-tags multiome with gene expression data from Slide-tags snRNA-seq using SelectIntegrationFeatures, FindIntegrationAnchors, and IntegrateData across all features with default parameters from Seurat (4.3.0). Integrated gene expression values were scaled and centred. For visualisation in two dimensions, we embedded nuclei in a Uniform Manifold Approximation and Projection (UMAP)^78^ using the top 30 PCs, with: number of neighbours =30, min_dist = 0.3, spread =1, local connectivity = 1, and the cosine distance metric. We identified shared nearest neighbours using the top 30 principal components. Clusters of similar cells were detected using the Louvain method for community detection, implemented using *FindClusters*, with a resolution = 1. Cells from Slide-tags multiome were annotated based on marker genes and co-clustering with Slide-tags snRNA-seq cells. Gene expression counts from Slide-tags multiome were re-scaled and re-cluster as described above using the non-integrated object for subsequent analyses.

#### 7.2 Inferring copy number variation

InferCNV (1.3.3) was used to infer large-scale copy number variation from standard snRNA-seq data and from snRNA-seq data from a 10x multiome experiment as previously recommended (inferCNV of the Trinity CTAT Project, https://github.com/broadinstitute/inferCNV). CellBender-corrected counts were extracted from annotated Seurat objects, where normal reference cells were specified as all cells not labelled as tumour. InferCNV was run under the following parameters: cutoff = 0.1, cluster_by_groups = T, denoise = T, HMM = T, num_threads = 60.

#### 7.3 T cell receptor analysis

TCR analyses focused on CD8 T cells where we used Fisher’s exact test to test if: (1) the beta chain sequence CASRASNEQFF was tumour compartment biassed compared against all CD8 T cells with profiled beta chains, where tumour compartment segmentation was performed manually based on tumour subpopulation density; and (2) paired CD8 T cells with TCR alpha chain CAEWYNQGGKLIF and beta chain CASRASNEQFF were tumour compartment biassed.

#### 7.4 ATAC analysis

Latent semantic indexing (LSI) was performed on the peaks assay using Signac, with the RunTFIDF and RunSVD functions. For visualisation in two dimensions, we embedded nuclei in a Uniform Manifold Approximation and Projection (UMAP)^78^ using LSI dimensions 2-30. Nuclei were visualised using the combination of modalities profiled, with weighted-nearest neighbour (WNN) analysis. Multimodal neighbours were identified using Seurat’s FindMultiModalNeighbors function, with the RNA PCA dimensions 1:50, and the ATAC LSI dimensions 2:50. These neighbours were then used as input into RunUMAP for visualisation.

In order to annotate the motifs present in peaks, the Signac function CreateMotifObject was used to create a motif object, with all human motifs from the Jaspar 2020 database. Motif accessibility z-scores were then calculated using Signac’s RunChromVAR function (ChromVAR 1.16.0). Gene activity scores were calculated using the Signac function GeneActivity. We normalised these gene scores by the total gene score per nucleus to the median nUMI for the RNA assay (NGS) and log-transformed these values to report gene expression as E = log(NGS + 1).

#### 7.5 Differential gene expression, differential chromatin gene scores, and gene set enrichment analysis

Differential gene expression analyses were performed using the MAST implemented in FindMarkers from Seurat^88^. Analysis comparing tumour cluster 1 and tumour cluster 2 from Slide-tags snRNA-seq and comparing compartment-specific CD8 T cells from Slide-tags multiome data used min.pct = 0.25 and log2fc.threshold = 0.25. Analysis comparing tumour cluster 1 and tumour cluster 2 from Slide-tags multiome data used min.pct = 0.1 and log2fc.threshold = 0.25. Gene ontology biological process (GO_Biological_Process_2021) gene set enrichment analysis was performed with the Enrichr package (3.1) in R^89–91^ on tumour cluster 2 enriched differentially expressed genes with log2FC < -0.5 and adjust *p* value < 0.05. Differential chromatin gene score analysis was conducted using the Wilcoxon Rank Sum test implemented in FindMarkers from Seurat with min.pct = 0.1 and log2fc.threshold = 0.

#### 7.6 Melanocytic-like and mesenchymal-like signatures

We scored tumour cells on melanocytic-like and mesenchymal-like signatures using AddModuleScore in Seurat with a list of genes adapted from previous work (Table S13)^50, 92^. Correlations of chromVar motif scores with mesenchymal scores were tested using Pearson’s correlation coefficient and *p* values were corrected using the Benjamini-Hochberg procedure. Spatial autocorrelations of chromVar motifs were tested using Moran.I from the ape package (5.6-2) in R^93^, where the weights matrix was specified as 1/distance^2^.

## Supplementary Material

### Supplementary Figures

**Supplementary Figure. 1:**
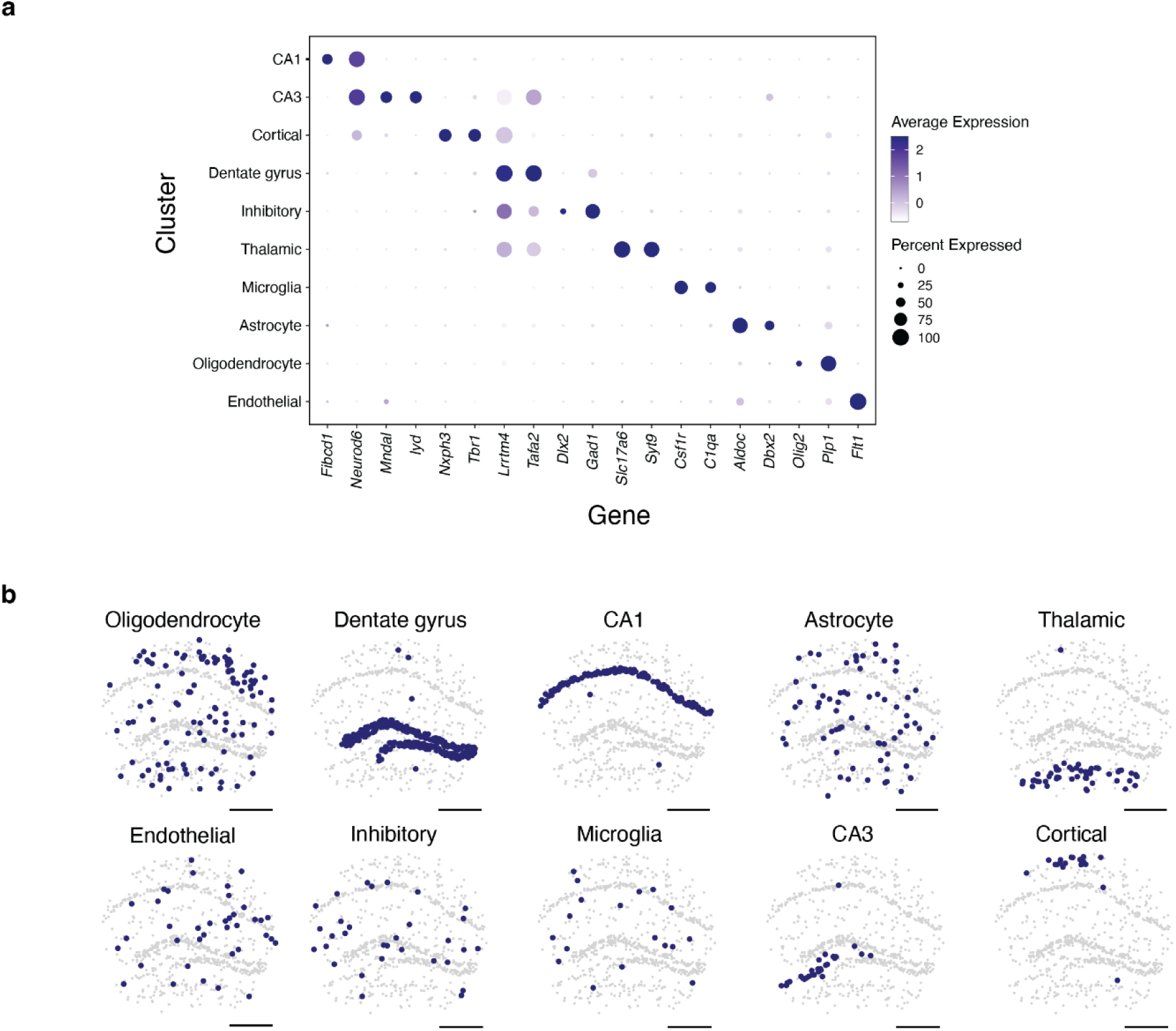
Cell type assignment and spatial mapping in the mouse hippocampus. **a,** Expression of marker genes by cell type cluster. **b,** Spatial positions of each cell by cell type cluster. All scale bars denote 500 μm. CA1 = Cornu Ammonis area 1, CA3 = Cornu Ammonis area 3.

**Supplementary Figure. 2:**
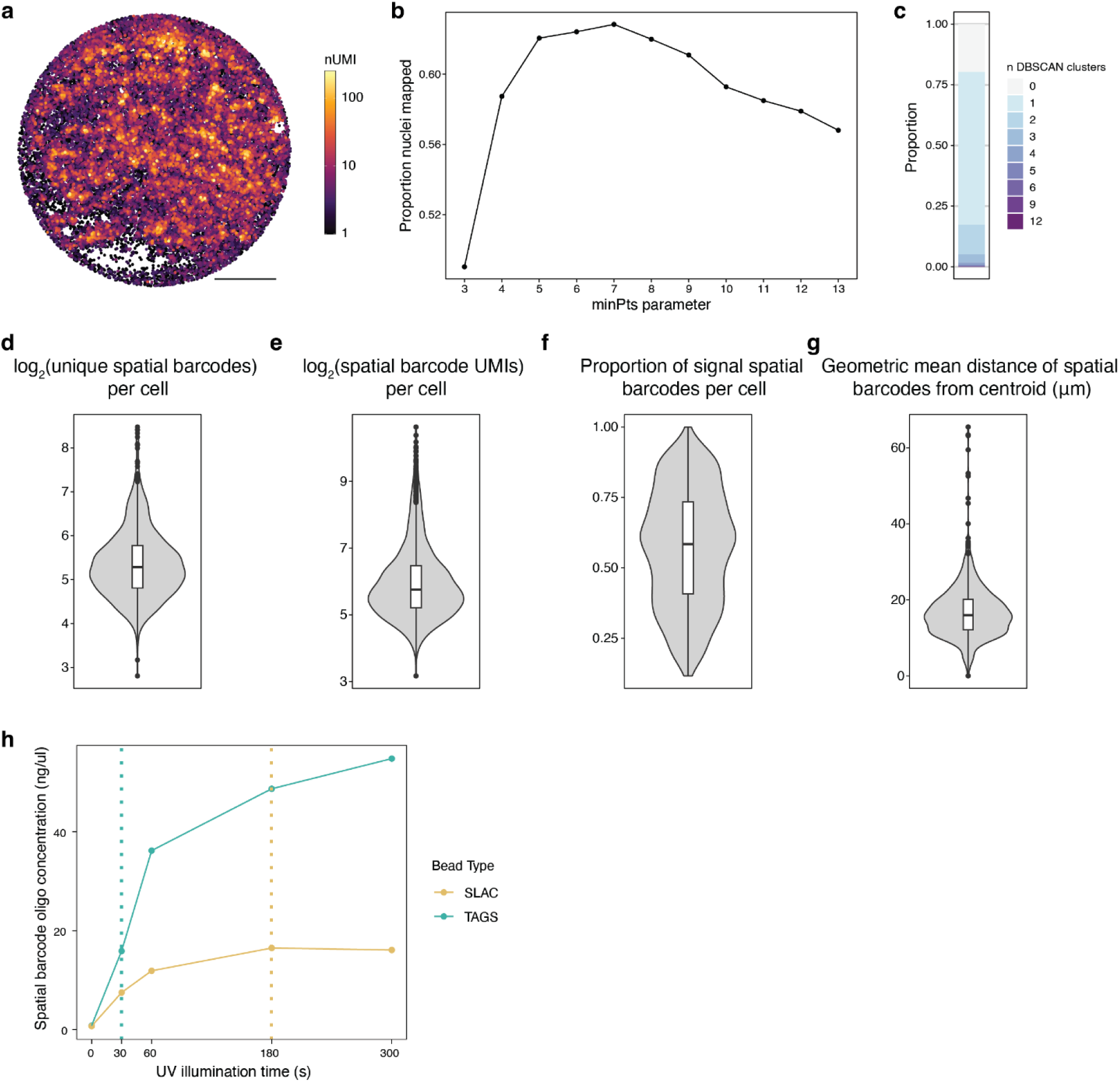
Assessing the mapping of single nuclei using spatial barcodes in the mouse hippocampus. **a,** Each recovered spatial barcode is shown coloured by the number of detected UMIs. **b,** The proportion of nuclei mapped for each minPts parameter tested in DBSCAN. **c,** The proportion of cells that are assigned to each number of DBSCAN clusters. **d-e,** Violin plots showing different spatial barcode metrics for every cell that is a spatial singlet. **f,** Violin plot showing the proportion of spatial barcode UMIs that are assigned to the DBSCAN singlet cluster (signal) vs. all other spatial barcode UMIs recovered for that cell. **g,** Violin plot showing the mean radial distance for spatial barcodes for each spatial singlet cluster. **h,** plot showing the concentration of oligos released by time under illumination at the same light source power, for each bead type used in Slide-tags experiments. The time used for cleavage for each bead type is shown with the dotted lines. Scale bar denotes 500 μm. Boxplots show: centre line, median; box limits, upper and lower quartiles; whiskers, 1.5x interquartile range; points, outliers.

**Supplementary Figure. 3:**
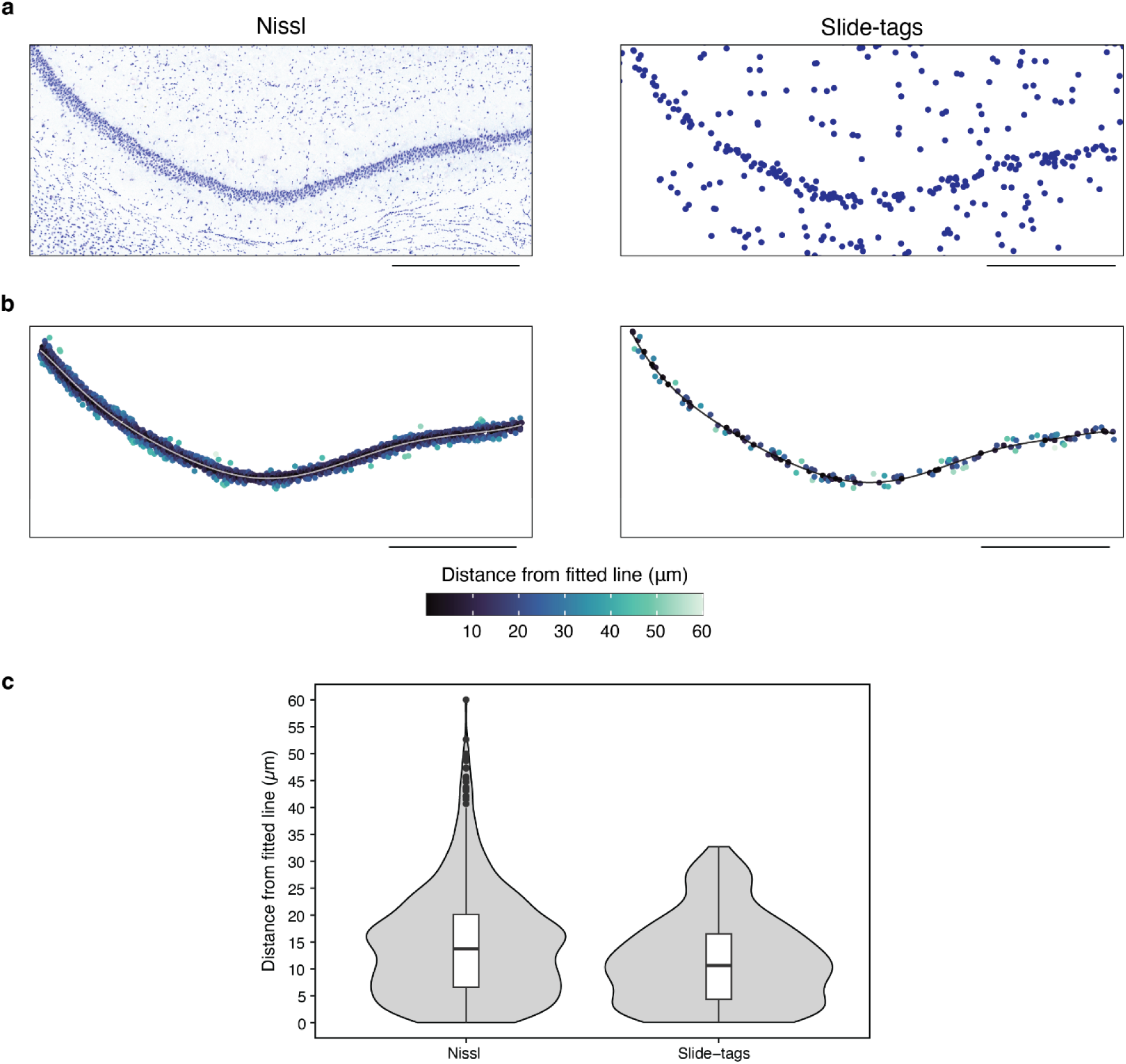
Spatial resolution measurements in the mouse hippocampus. **a,** A 10 um nissl-stained section (left) was taken adjacently to the Slide-tags profiled section (right). **b,** The CA1 cells were subsetted in each case and a line was fitted to measure the midpoint of this structure. Orthogonal distances from this midpoint were then calculated and points are coloured by this distance. **c,** Violin plots showing the distribution of distances from the fitted line in b. Boxplots show: centre line, median; box limits, upper and lower quartiles; whiskers, 1.5x interquartile range; points, outliers. All scale bars denote 500 μm.

**Supplementary Figure. 4:**
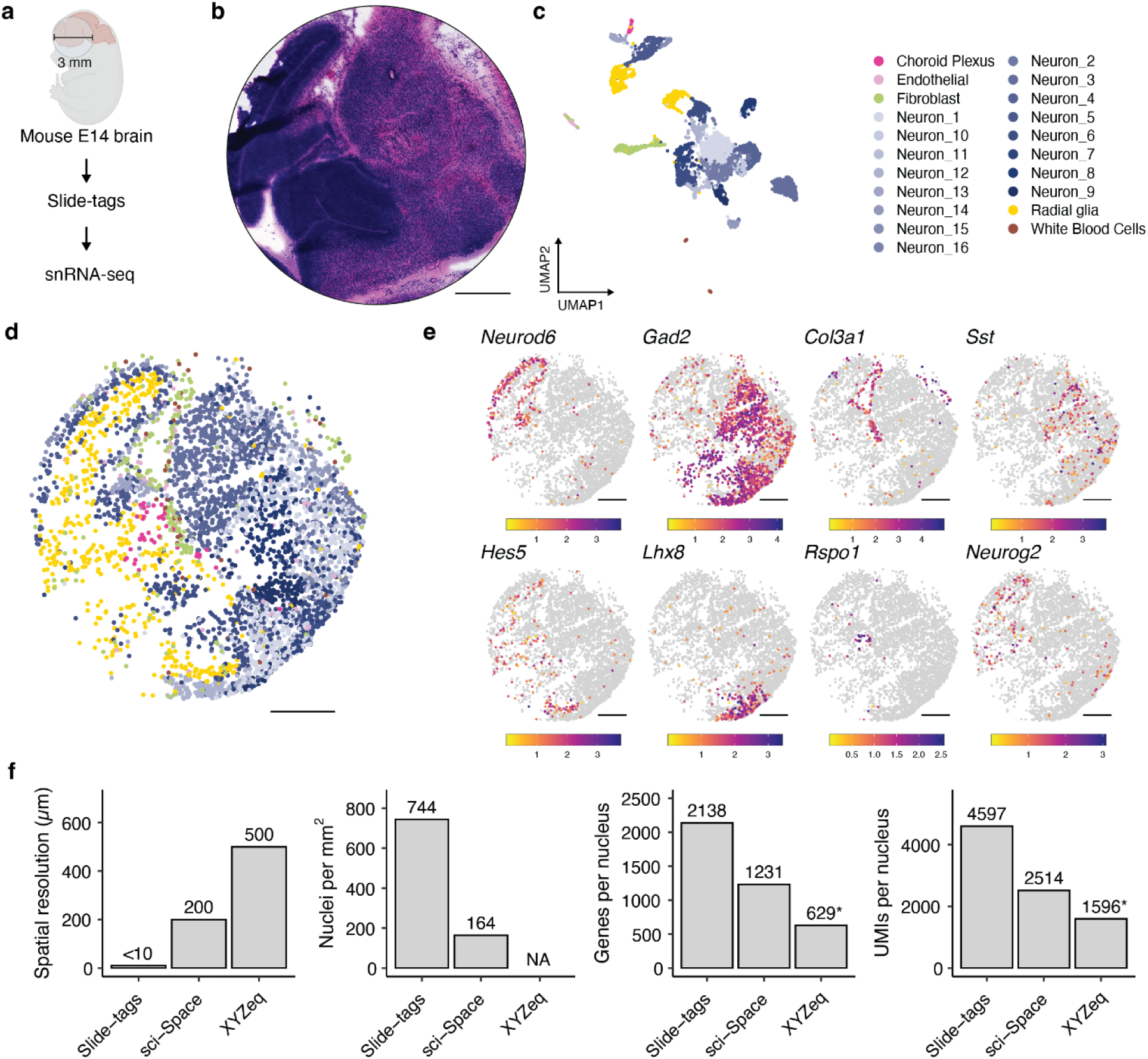
Slide-tags snRNA-seq applied to the embryonic mouse brain at E14. **a,** Schematic of Slide-tags snRNA-seq on a 3 mm diameter region of the embryonic mouse brain at E14. **b.** A haematoxylin and eosin stained section which was adjacent to the profiled section. **c.** UMAP embedding of snRNA-seq profiles coloured by cell-type annotations. **d.** Spatial positions of cells coloured as in C. **e.** Spatial marker gene expression. Expression counts for each cell were divided by the total counts for that cell and multiplied by 10,000, this value + 1 is then natural-log transformed. **f.** Comparison metrics plotted for Slide-tags snRNA-seq on the mouse E14 embryonic brain. * = XYZeq was not performed on embryonic mouse brain at E14 and so these metrics may not be directly comparable due to tissue-specific effects. All scale bars denote 500 μm.

**Supplementary Figure 5.**
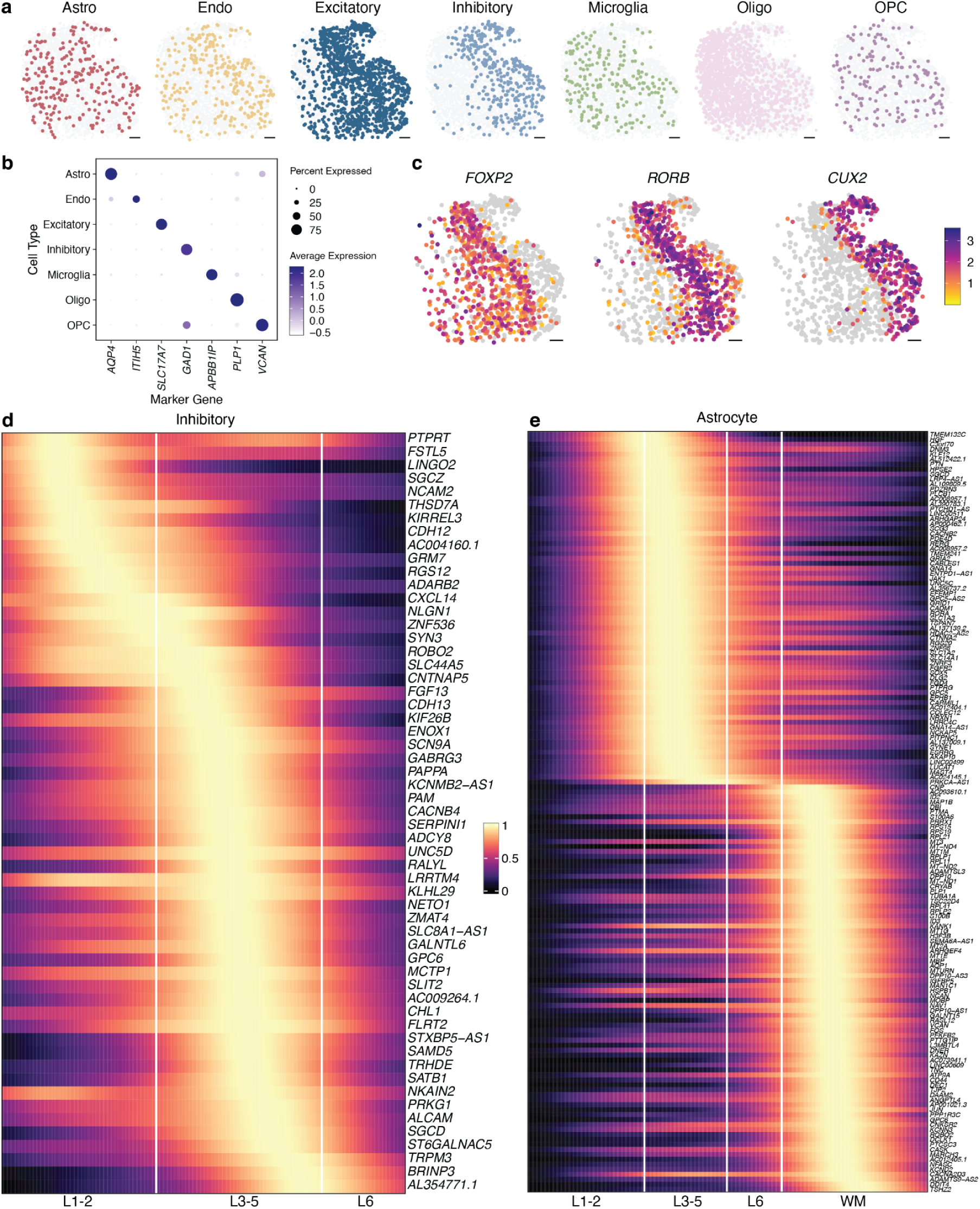
Slide-tags snRNA-seq applied to the human brain enables spatial mapping of cell types and cell-type specific spatially varying gene expression. **a,** Individual plots of per-cell type spatial distribution. **b,** Dotplot showing the marker genes used to assign cell types to clusters. **c,** The gene expression distribution of three canonical layer marker genes in excitatory neurons. **d,e,** A 1D gene expression heatmap for genes in inhibitory neurons and astrocytes. All scale bars represent 500 µm. Oligo = Oligodendrocyte, OPC = Oligodendrocyte precursor cell, Astro = Astrocyte, INH = Inhibitory, EX = Excitatory, GM = Grey matter, WM = White matter. Gene names and details in Supplementary Table 1.

**Supplementary Figure 6.**
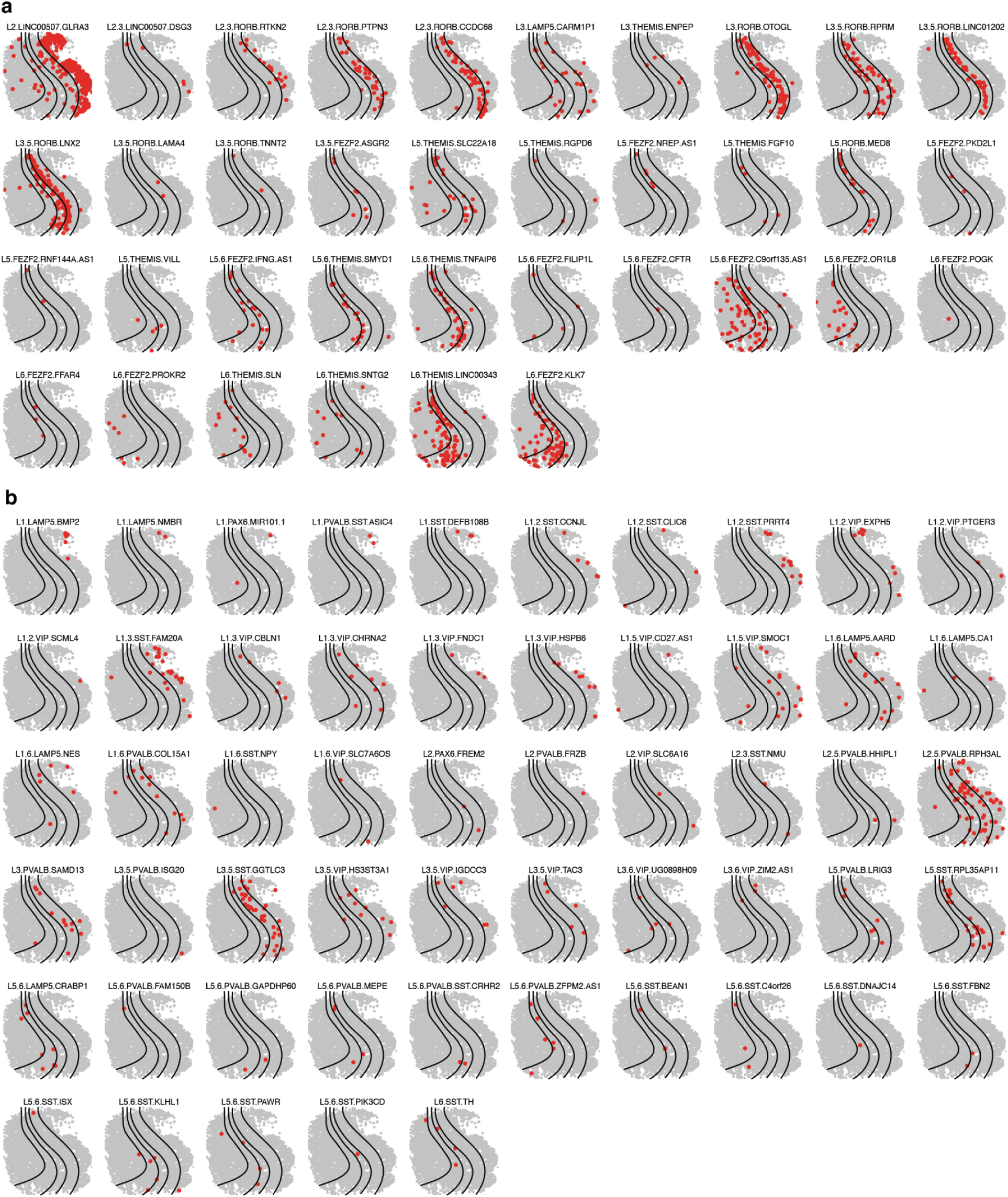
Slide-tags snRNA-seq in the human brain enables mapping of neuron sub-cell-types. **a,** Excitatory neuron subtypes plotted by spatial location. **b,** Inhibitory neuron subtypes plotted by spatial location. Subtype names from Bakken et al., 2021.

**Supplementary Figure 7.**
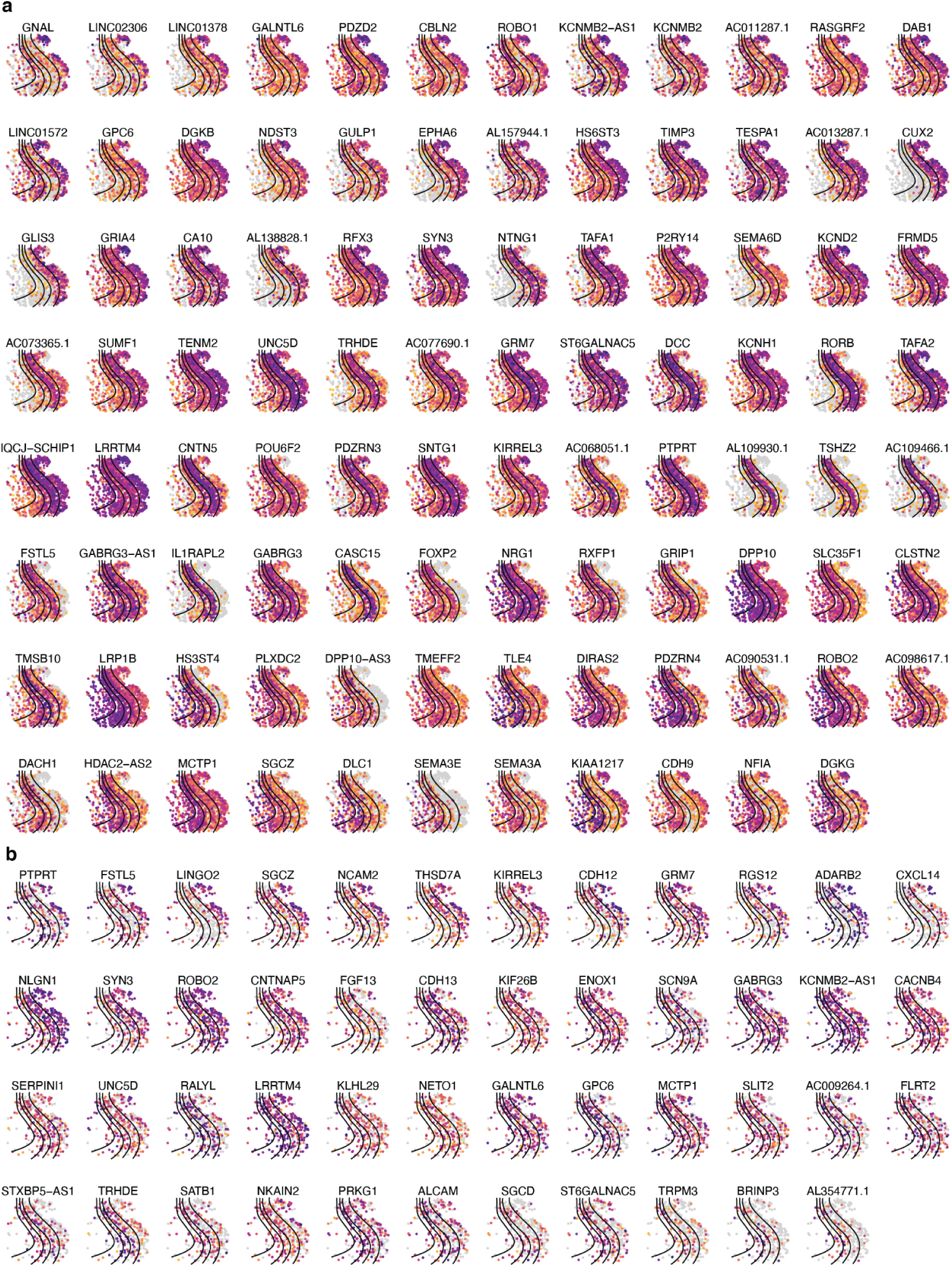
Slide-tags snRNA-seq in the human brain reveals cell-type specific spatial gradients of gene expression in neurons. Spatially varying genes identified in: **a,** Excitatory neurons, **b,** Inhibitory neurons

**Supplementary Figure 8.**
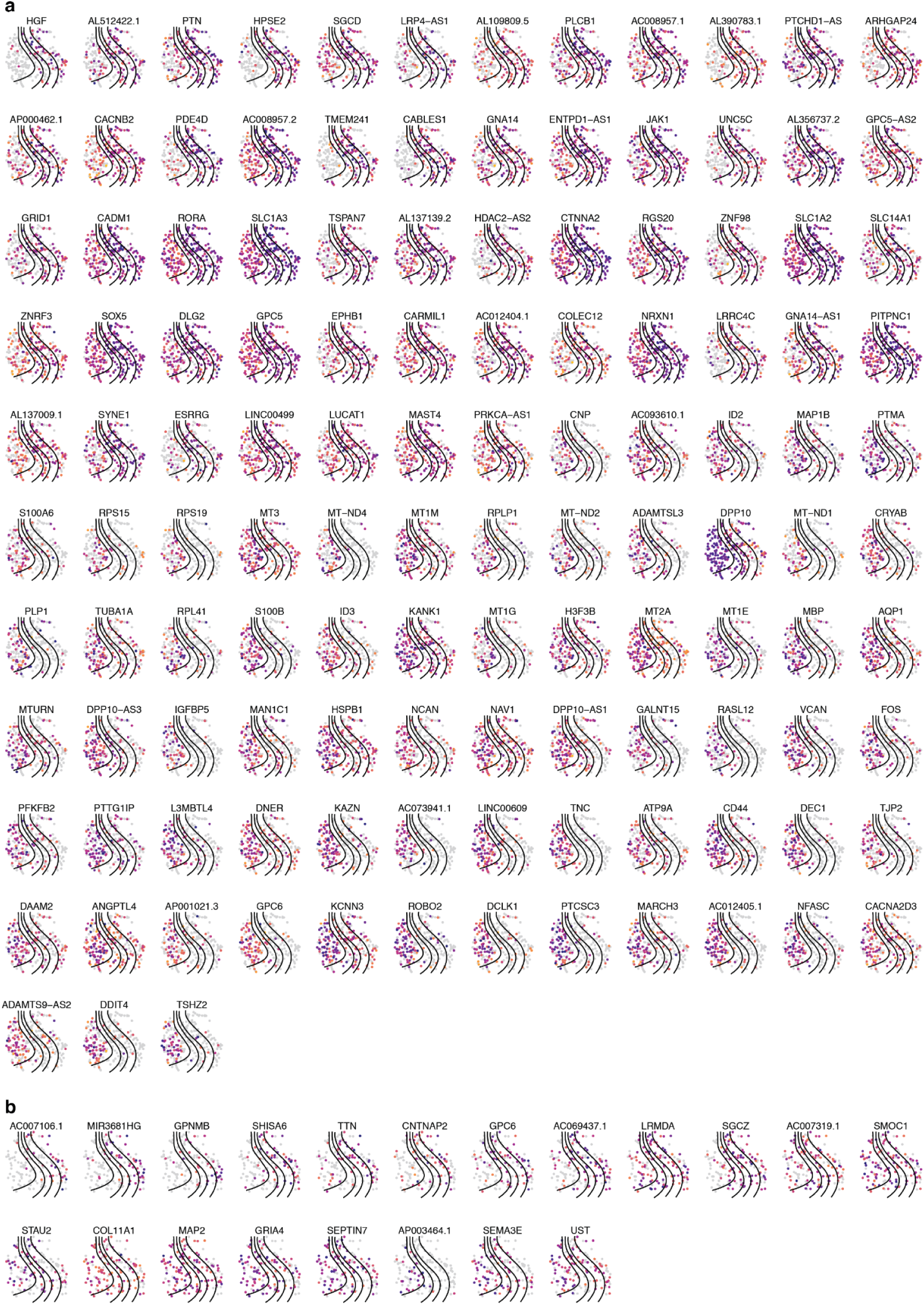
Slide-tags snRNA-seq in the human brain reveals cell-type specific spatial gradients of gene expression in non-neuronal cell types. Spatially varying genes identified in: **a,** Astrocytes, **b,** Oligodendrocyte Precursor Cells (OPCs).

**Supplementary Figure 9.**
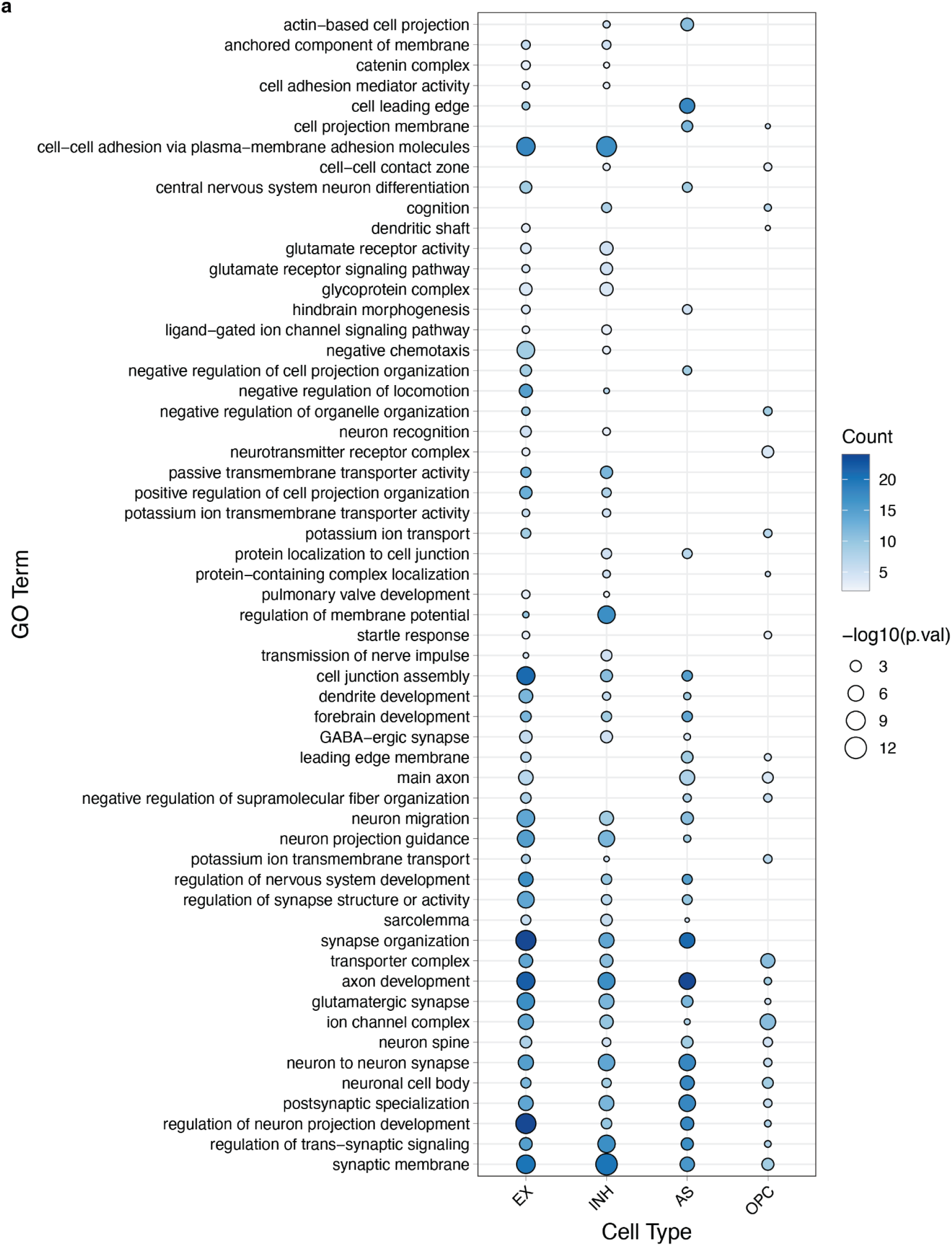
**Gene Ontology Enrichment of cell-type specific spatial gradient genes revealed by Slide-tags snRNA-seq in the human brain.**

**Supplementary Figure 10.**
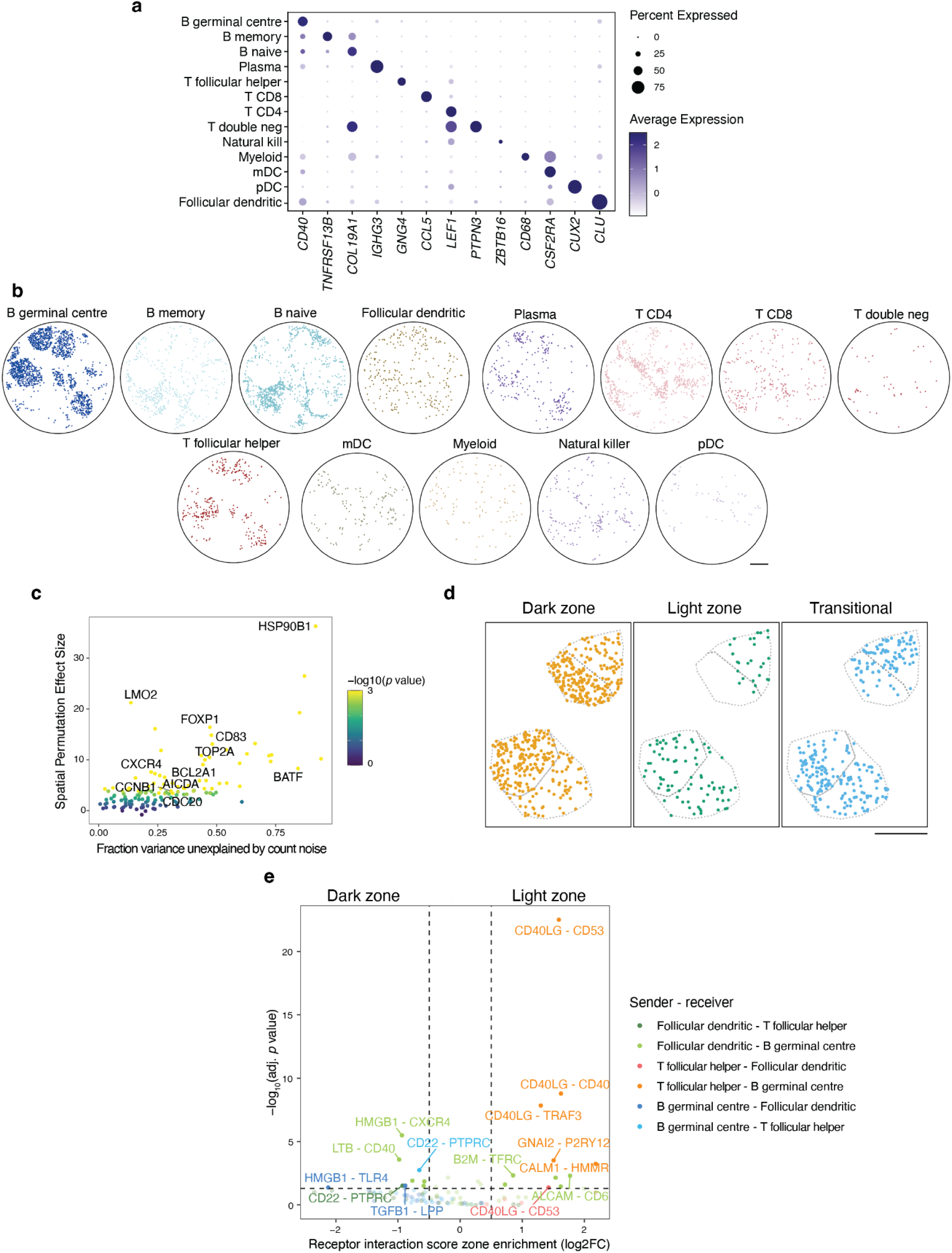
Receptor-ligand prediction from Slide-tags human tonsil data. **a,** Expression of select marker genes by cell type cluster. **b,** Spatial mapping of cell types. **c,** Scatter plot of gene expression variance not explained by count noise and spatial permutation effect size of previously reported dark zone and light zone marker genes. **d,** Spatial mapping of dark zone, light zone, and transitional germinal centre B cells in two representative germinal centres. **e,** Volcano plot of receptor interaction intensity scores compared between zones in two representative germinal centres. All scale bars denote 500 μm. T double neg = T double negative, mDC = myeloid dendritic cells, pDC = plasmacytoid dendritic cells.

**Supplementary Figure 11.**
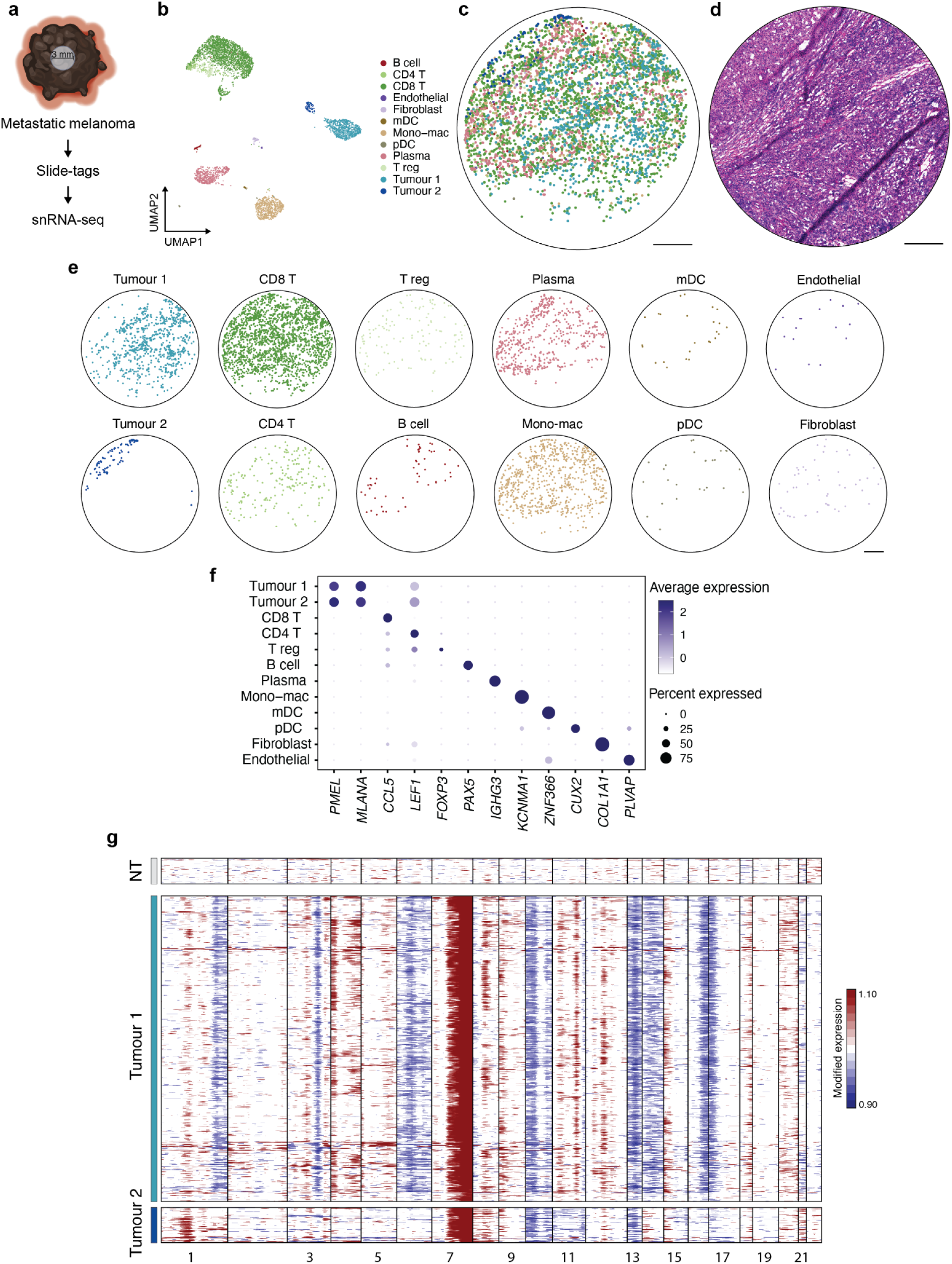
Slide-tags snRNA-seq on human melanoma. **a,** Schematic representation of Slide-tags snRNA-seq on a 3 mm circular region of a human melanoma lymph node metastasis. **b,** UMAP embeddings of snRNA-seq profiles coloured by cell type. **c,** Spatial mapping of cell types. **d,** Adjacent H&E-stained section of the profiled region. **e,** Spatial mapping of profiled cell types. **f,** Expression of select marker genes by cell type cluster. **g,** Inferred copy number alterations from transcriptomic data. NT indicates a representative subset of non-tumour cells. All scale bars denote 500 μm. T reg = T regulatory cells, mDC = myeloid dendritic cells, Mono-mac = monocyte-derived macrophages, pDC = plasmacytoid dendritic cells.

**Supplementary Figure 12.**
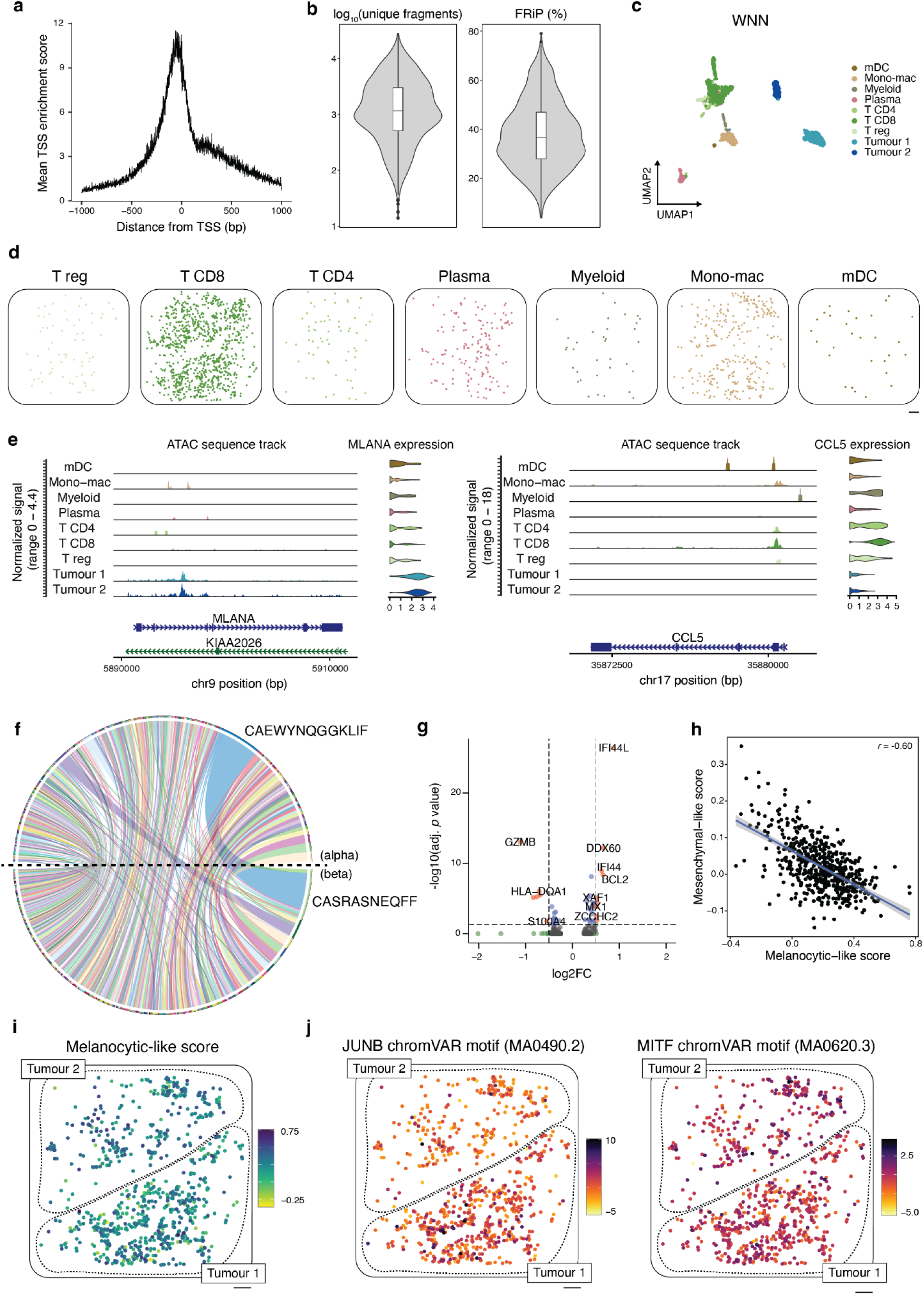
Slide-tags multiome on human melanoma. **a,** Mean TSS enrichment score. **b,** Violin plots of log_10_-transformed unique fragments and fraction of reads in peaks (FRiP) percentage. Boxplots show: centre line, median; box limits, upper and lower quartiles; whiskers, 1.5x interquartile range; points, outliers. **c,** Weighted nearest neighbour UMAP embeddings of snRNA-seq and snATAC-seq profiles coloured by cell type. **d,** Spatial mapping of cell types. **e,** ATAC sequence track and gene expression violin plot of *MLANA* and *CCL5* across cell types. **f,** TCR pairing chord plot of alpha and beta chain pairing frequencies in CD8 T cells. **g,** Differential gene expression volcano plot between CD8 T cells in tumour compartment 1 vs tumour compartment 2. **h,** Scatter plot of melanocytic-like scores and mesenchymal-like scores of tumour cluster 1 cells in tumour compartment 1. Pearson’s *r* value is reported. **i,** Mesenchymal-like cell state score spatial distribution. **j,** Spatial distribution of JUNB and MITF chromVAR motif scores. All scale bars denote 500 μm. T reg = T regulatory cells, mDC = myeloid dendritic cells, Mono-mac = monocyte-derived macrophages.

**Supplementary Figure 13.**
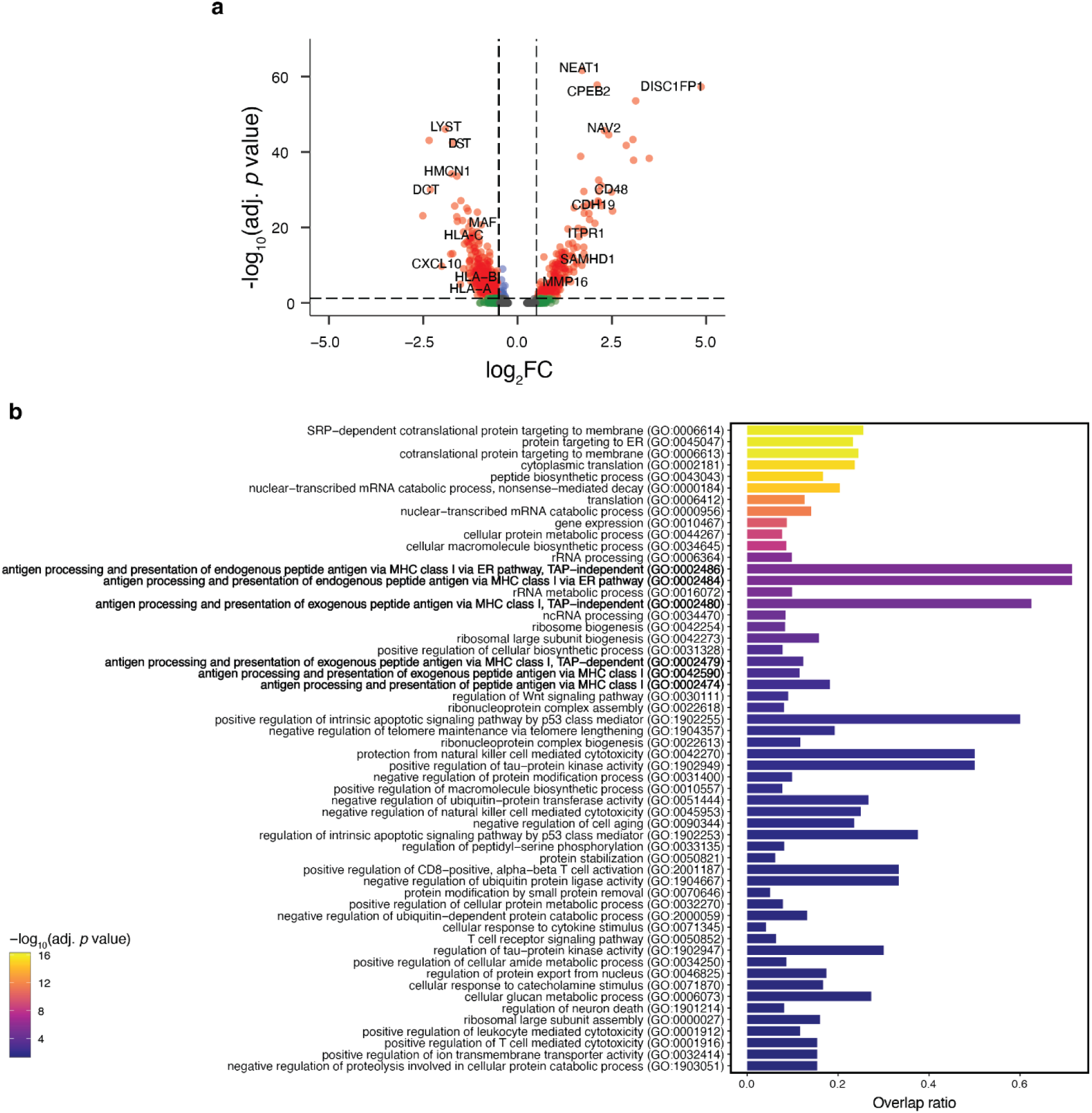
Differential gene expression and gene set enrichment analysis between tumour cluster 1 and 2. **a,** Volcano plot of differentially expressed genes comparing tumour cluster 1 against tumour cluster 2 from the Slide-tags snRNA-seq run. **b,** Gene ontology biological process (GO_Biological_Process_2021) gene set enrichment analysis on genes upregulated in tumour cluster 2 (negative log_2_FC) compared with tumour cluster 1 from the Slide-tags snRNA-seq experiment.

### Supplementary Tables

**Supplementary Table 1.** Per-celltype gene list ranked by spatial variation, human cortex

**Supplementary Table 2.** GO enrichment analysis results, human cortex

**Supplementary Table 3.** Spatial varying genes in germinal centres, human tonsil.

**Supplementary Table 4.** Spatial effect size from spatial varying gene expression results and percent variance in gene expression, human tonsil.

**Supplementary Table 5.** Receptor-ligand interaction prediction results, human tonsil.

**Supplementary Table 6.** Germinal centre zone enrichment test of receptor-ligand interactions, human tonsil.

**Supplementary Table 7.** Compartment-specific T cell receptor sequences of CD8 T cells, human melanoma.

**Supplementary Table 8.** Differential gene expression between CD8 T cells in tumour compartment 1 and tumour compartment 2 from Slide-tags multiome performed on human melanoma.

**Supplementary Table 9.** Differential gene expression between tumour cluster 1 and tumour cluster 2 from Slide-tags snRNA-seq performed on human melanoma.

**Supplementary Table 10.** Differential gene expression (x) and differential chromatin gene scores (y) between tumour cluster 1 and tumour cluster 2 from Slide-tags multiome performed on human melanoma.

**Supplementary Table 11.** Spatial autocorrelation of ChromVAR transcription factor motifs correlated with mesenchymal-like scores in tumour cluster 1, human melanoma.

**Supplementary Table 12.** Metadata and data pre-processing details for each Slide-tags experiment.

**Supplementary Table 13.** Mesenchymal-like and melanocytic-like genes used to score tumour cells, human melanoma.

## References

1. Stoeckius, M. et al. Simultaneous epitope and transcriptome measurement in single cells. Nat. Methods 14, 865–868 (2017).

2. Zheng, G. X. Y. et al. Massively parallel digital transcriptional profiling of single cells. Nat. Commun. 8, 14049 (2017).

3. Macosko, E. Z. et al. Highly Parallel Genome-wide Expression Profiling of Individual Cells Using Nanoliter Droplets. Cell 161, 1202–1214 (2015).

4. Cusanovich, D. A. et al. Multiplex single cell profiling of chromatin accessibility by combinatorial cellular indexing. Science 348, 910–914 (2015).

5. Klein, A. M. et al. Droplet barcoding for single-cell transcriptomics applied to embryonic stem cells. Cell 161, 1187–1201 (2015).

6. Vitak, S. A. et al. Sequencing thousands of single-cell genomes with combinatorial indexing. Nat. Methods 14, 302–308 (2017).

7. Mulqueen, R. M. et al. Highly scalable generation of DNA methylation profiles in single cells. Nat. Biotechnol. 36, 428–431 (2018).

8. Zhao, T. et al. Spatial genomics enables multi-modal study of clonal heterogeneity in tissues. Nature 601, 85–91 (2022).

9. Ståhl, P. L. et al. Visualization and analysis of gene expression in tissue sections by spatial transcriptomics. Science 353, 78–82 (2016).

10. Rodriques, S. G. et al. Slide-seq: A scalable technology for measuring genome-wide expression at high spatial resolution. Science 363, 1463–1467 (2019).

11. Liu, Y. et al. High-Spatial-Resolution Multi-Omics Sequencing via Deterministic Barcoding in Tissue. Cell 183, 1665–1681.e18 (2020).

12. Nichterwitz, S. et al. Laser capture microscopy coupled with Smart-seq2 for precise spatial transcriptomic profiling. Nat. Commun. 7, 12139 (2016).

13. Deng, Y. et al. Spatial profiling of chromatin accessibility in mouse and human tissues. Nature 609, 375–383 (2022).

14. Llorens-Bobadilla, E. et al. Solid-phase capture and profiling of open chromatin by spatial ATAC. Nat. Biotechnol. (2023) doi:10.1038/s41587-022-01603-9.

15. Zhang, D. et al. Spatial epigenome-transcriptome co-profiling of mammalian tissues. Nature (2023) doi:10.1038/s41586-023-05795-1.

16. Palla, G., Fischer, D. S., Regev, A. & Theis, F. J. Spatial components of molecular tissue biology. Nat. Biotechnol. (2022) doi:10.1038/s41587-021-01182-1.

17. Cable, D. M. et al. Robust decomposition of cell type mixtures in spatial transcriptomics. Nat. Biotechnol. 40, 517–526 (2022).

18. Kleshchevnikov, V. et al. Cell2location maps fine-grained cell types in spatial transcriptomics. Nat. Biotechnol. 40, 661–671 (2022).

19. Biancalani, T. et al. Deep learning and alignment of spatially resolved single-cell transcriptomes with Tangram. Nat. Methods 18, 1352–1362 (2021).

20. Srivatsan, S. R. et al. Embryo-scale, single-cell spatial transcriptomics. Science 373, 111–117 (2021).

21. Lee, Y. et al. XYZeq: Spatially resolved single-cell RNA sequencing reveals expression heterogeneity in the tumor microenvironment. Sci Adv 7, (2021).

22. Stickels, R. R. et al. Highly sensitive spatial transcriptomics at near-cellular resolution with Slide-seqV2. Nature Biotechnology vol. 39 313–319 Preprint at https://doi.org/10.1038/s41587-020-0739-1 (2021).

23. Hao, Y. et al. Integrated analysis of multimodal single-cell data. Cell 184, 3573–3587.e29 (2021).

24. Xu, X., Ester, M., Kriegel, H.-P. & Sander, J. A distribution-based clustering algorithm for mining in large spatial databases. Proceedings 14th International Conference on Data Engineering Preprint at https://doi.org/10.1109/icde.1998.655795.

25. Lein, E. S. et al. Genome-wide atlas of gene expression in the adult mouse brain. Nature 445, 168–176 (2007).

26. Chen, A. et al. Spatiotemporal transcriptomic atlas of mouse organogenesis using DNA nanoball-patterned arrays. Cell 185, 1777–1792.e21 (2022).

27. Maynard, K. R. et al. Transcriptome-scale spatial gene expression in the human dorsolateral prefrontal cortex. Nat. Neurosci. 24, 425–436 (2021).

28. Bakken, T. E. et al. Comparative cellular analysis of motor cortex in human, marmoset and mouse. Nature 598, 111–119 (2021).

29. Victora, G. D. & Nussenzweig, M. C. Germinal Centers. Annu. Rev. Immunol. 40, 413–442 (2022).

30. Allen, C. D. C. et al. Germinal center dark and light zone organization is mediated by CXCR4 and CXCR5. Nat. Immunol. 5, 943–952 (2004).

31. Revy, P. et al. Activation-induced cytidine deaminase (AID) deficiency causes the autosomal recessive form of the Hyper-IgM syndrome (HIGM2). Cell 102, 565–575 (2000).

32. Muramatsu, M. et al. Class switch recombination and hypermutation require activation-induced cytidine deaminase (AID), a potential RNA editing enzyme. Cell 102, 553–563 (2000).

33. Ottina, E. et al. Targeting antiapoptotic A1/Bfl-1 by in vivo RNAi reveals multiple roles in leukocyte development in mice. Blood 119, 6032–6042 (2012).

34. Osada, H., Grutz, G., Axelson, H., Forster, A. & Rabbitts, T. H. Association of erythroid transcription factors: complexes involving the LIM protein RBTN2 and the zinc-finger protein GATA1. Proc. Natl. Acad. Sci. U. S. A. 92, 9585–9589 (1995).

35. Wadman, I. A. et al. The LIM-only protein Lmo2 is a bridging molecule assembling an erythroid, DNA-binding complex which includes the TAL1, E47, GATA-1 and Ldb1/NLI proteins. EMBO J. 16, 3145–3157 (1997).

36. Papa, I. & Vinuesa, C. G. Synaptic Interactions in Germinal Centers. Front. Immunol. 9, 1858 (2018).

37. Dimitrov, D. et al. Comparison of methods and resources for cell-cell communication inference from single-cell RNA-Seq data. Nat. Commun. 13, 3224 (2022).

38. Elgueta, R. et al. Molecular mechanism and function of CD40/CD40L engagement in the immune system. Immunol. Rev. 229, 152–172 (2009).

39. Ni, C. Z. et al. Molecular basis for CD40 signaling mediated by TRAF3. Proc. Natl. Acad. Sci. U. S. A. 97, 10395–10399 (2000).

40. Tam, W. L. & Weinberg, R. A. The epigenetics of epithelial-mesenchymal plasticity in cancer. Nat. Med. 19, 1438–1449 (2013).

41. Liau, B. B. et al. Adaptive Chromatin Remodeling Drives Glioblastoma Stem Cell Plasticity and Drug Tolerance. Cell Stem Cell 20, 233–246.e7 (2017).

42. Sharma, S. V. et al. A chromatin-mediated reversible drug-tolerant state in cancer cell subpopulations. Cell 141, 69–80 (2010).

43. Suvà, M. L., Riggi, N. & Bernstein, B. E. Epigenetic reprogramming in cancer. Science 339, 1567–1570 (2013).

44. Lomakin, A. et al. Spatial genomics maps the structure, nature and evolution of cancer clones. Nature 611, 594–602 (2022).

45. Erickson, A. et al. Spatially resolved clonal copy number alterations in benign and malignant tissue. Nature 608, 360–367 (2022).

46. Gerstung, M. et al. The evolutionary history of 2,658 cancers. Nature 578, 122–128 (2020).

47. Balázs, M. et al. Chromosomal imbalances in primary and metastatic melanomas revealed by comparative genomic hybridization. Cytometry 46, 222–232 (2001).

48. Tickle, T., Tirosh, I., Georgescu, C., Brown, M. & Haas, B. inferCNV of the Trinity CTAT Project. (2019).

49. Liu, S. et al. Spatial maps of T cell receptors and transcriptomes reveal distinct immune niches and interactions in the adaptive immune response. Immunity 55, 1940–1952.e5 (2022).

50. Berico, P., et al. CDK7 and MITF repress a transcription program involved in survival and drug tolerance in melanoma. EMBO Rep. 22, e51683 (2021).

51. Shao, H., Kirkwood, J. M. & Wells, A. Tenascin-C Signaling in melanoma. Cell Adh. Migr. 9, 125–130 (2015).

52. Fukunaga-Kalabis, M. et al. Tenascin-C promotes melanoma progression by maintaining the ABCB5-positive side population. Oncogene 29, 6115–6124 (2010).

53. Schep, A. N., Wu, B., Buenrostro, J. D. & Greenleaf, W. J. chromVAR: inferring transcription-factor-associated accessibility from single-cell epigenomic data. Nat. Methods 14, 975–978 (2017).

54. Pedri, D., Karras, P., Landeloos, E., Marine, J.-C. & Rambow, F. Epithelial-to-mesenchymal-like transition events in melanoma. FEBS J. 289, 1352–1368 (2022).

55. Tirosh, I. et al. Dissecting the multicellular ecosystem of metastatic melanoma by single-cell RNA-seq. Science 352, 189–196 (2016).

56. Shaffer, S. M., et al. Memory Sequencing Reveals Heritable Single-Cell Gene Expression Programs Associated with Distinct Cellular Behaviors. Cell 182, 947–959.e17 (2020).

57. Shaffer, S. M., et al. Rare cell variability and drug-induced reprogramming as a mode of cancer drug resistance. Nature 546, 431–435 (2017).

58. Navin, N. et al. Tumour evolution inferred by single-cell sequencing. Nature 472, 90–94 (2011).

59. Gonzalez-Pena, V. et al. Accurate genomic variant detection in single cells with primary template-directed amplification. Proc. Natl. Acad. Sci. U. S. A. 118, (2021).

60. Farlik, M. et al. Single-cell DNA methylome sequencing and bioinformatic inference of epigenomic cell-state dynamics. Cell Rep. 10, 1386–1397 (2015).

61. Smallwood, S. A. et al. Single-cell genome-wide bisulfite sequencing for assessing epigenetic heterogeneity. Nat. Methods 11, 817–820 (2014).

62. Bartosovic, M., Kabbe, M. & Castelo-Branco, G. Single-cell CUT&Tag profiles histone modifications and transcription factors in complex tissues. Nat. Biotechnol. 39, 825–835 (2021).

63. Kaya-Okur, H. S., et al. CUT&Tag for efficient epigenomic profiling of small samples and single cells. Nat. Commun. 10, 1930 (2019).

64. Chung, H. et al. Joint single-cell measurements of nuclear proteins and RNA in vivo. Nat. Methods 18, 1204–1212 (2021).

65. Chen, A. F. et al. NEAT-seq: simultaneous profiling of intra-nuclear proteins, chromatin accessibility and gene expression in single cells. Nat. Methods 19, 547–553 (2022).

66. Granja, J. M. et al. ArchR is a scalable software package for integrative single-cell chromatin accessibility analysis. Nat. Genet. 53, 403–411 (2021).

67. Clark, I. C. et al. Microfluidics-free single-cell genomics with templated emulsification. Nat. Biotechnol. (2023) doi:10.1038/s41587-023-01685-z.

68. Miller, T. E. et al. Mitochondrial variant enrichment from high-throughput single-cell RNA sequencing resolves clonal populations. Nat. Biotechnol. 40, 1030–1034 (2022).

69. Ludwig, L. S. et al. Lineage Tracing in Humans Enabled by Mitochondrial Mutations and Single-Cell Genomics. Cell 176, 1325–1339.e22 (2019).

70. La Manno, G. et al. RNA velocity of single cells. Nature 560, 494–498 (2018).

71. Slyper, M. et al. A single-cell and single-nucleus RNA-Seq toolbox for fresh and frozen human tumors. Nat. Med. 26, 792–802 (2020).

72. Fleming, S. J. et al. Unsupervised removal of systematic background noise from droplet-based single-cell experiments using CellBender. bioRxiv 791699 (2022) doi:10.1101/791699.

73. Ester, M., Kriegel, H.-P., Sander, J. & Xu, X. A density-based algorithm for discovering clusters in large spatial databases with noise. in Proceedings of the Second International Conference on Knowledge Discovery and Data Mining 226–231 (AAAI Press, 1996).

74. Hahsler, M., Piekenbrock, M. & Doran, D. dbscan: Fast Density-Based Clustering with R. J. Stat. Softw. 91, 1–30 (2019).

75. Bolotin, D. A. et al. MiXCR: software for comprehensive adaptive immunity profiling. Nat. Methods 12, 380–381 (2015).

76. Bolotin, D. A. et al. Antigen receptor repertoire profiling from RNA-seq data. Nat. Biotechnol. 35, 908–911 (2017).

77. Stuart, T. et al. Comprehensive Integration of Single-Cell Data. Cell vol. 177 1888–1902.e21 Preprint at https://doi.org/10.1016/j.cell.2019.05.031 (2019).

78. McInnes, L., Healy, J. & Melville, J. UMAP: Uniform Manifold Approximation and Projection for Dimension Reduction. arXiv [stat.ML] (2018).

79. Massoni-Badosa, R. et al. An Atlas of Cells in the Human Tonsil. Preprint at https://doi.org/10.1101/2022.06.24.497299.

80. Victora, G. D. et al. Identification of human germinal center light and dark zone cells and their relationship to human B-cell lymphomas. Blood 120, 2240–2248 (2012).

81. Chen, Y. A new methodology of spatial cross-correlation analysis. PLoS One 10, e0126158 (2015).

82. Choudhary, S. & Satija, R. Comparison and evaluation of statistical error models for scRNA-seq. Genome Biol. 23, 27 (2022).

83. Anselin, L. Local indicators of spatial association-LISA. Geogr. Anal. 27, 93–115 (2010).

84. Stuart, T., Srivastava, A., Madad, S., Lareau, C. A. & Satija, R. Single-cell chromatin state analysis with Signac. Nat. Methods 18, 1333–1341 (2021).

85. Zhang, Y. et al. Model-based analysis of ChIP-Seq (MACS). Genome Biol. 9, R137 (2008).

86. Arora, S., Morgan, M., Carlson, M. & Pagès, H. GenomeInfoDb: Utilities for manipulating chromosome names, including modifying them to follow a particular naming style. Preprint at https://bioconductor.org/packages/GenomeInfoDb (2023).

87. Amemiya, H. M., Kundaje, A. & Boyle, A. P. The ENCODE Blacklist: Identification of Problematic Regions of the Genome. Sci. Rep. 9, 9354 (2019).

88. Finak, G. et al. MAST: a flexible statistical framework for assessing transcriptional changes and characterizing heterogeneity in single-cell RNA sequencing data. Genome Biol. 16, 278 (2015).

89. Chen, E. Y. et al. Enrichr: interactive and collaborative HTML5 gene list enrichment analysis tool. BMC Bioinformatics 14, 128 (2013).

90. Kuleshov, M. V. et al. Enrichr: a comprehensive gene set enrichment analysis web server 2016 update. Nucleic Acids Res. 44, W90–7 (2016).

91. Xie, Z. et al. Gene Set Knowledge Discovery with Enrichr. Curr Protoc 1, e90 (2021).

92. Widmer, D. S. et al. Systematic classification of melanoma cells by phenotype-specific gene expression mapping. Pigment Cell Melanoma Res. 25, 343–353 (2012).

93. Paradis, E. & Schliep, K. ape 5.0: an environment for modern phylogenetics and evolutionary analyses in R. Bioinformatics 35, 526–528 (2019).

